# Systematic disruption of zebrafish fibrillin genes identifies a translational zebrafish model for Marfan syndrome

**DOI:** 10.1101/2025.06.21.659830

**Authors:** Karo De Rycke, Marina Horvat, Lisa Caboor, Petra Vermassen, Griet De Smet, Marta Santana Silva, Wouter Steyaert, Matthias Van Impe, Patrick Segers, Julie De Backer, Patrick Sips

## Abstract

**Background:** Fibrillins are essential components of the extracellular matrix. Marfan syndrome (MFS), the most common fibrillinopathy, is characterized by severe cardiovascular complications, including cardiac valve abnormalities, myocardial dysfunction, arrhythmias, and, most commonly, thoracic aortic disease. Unfortunately, no definitive medical cure is available.

**Objectives:** To establish a zebrafish model of MFS, to enhance understanding of the cardiovascular consequences of fibrillin impairment and identify novel therapeutic targets.

**Methods:** CRISPR/Cas9 technology was used to systematically target all zebrafish fibrillin genes. The cardiovascular phenotype was investigated using fluorescent microscopy at embryonic stages and cardiac ultrasound, histology, and synchrotron X-ray imaging in adults. RNA sequencing and drug testing were performed during early development.

**Results:** Fibrillin-2b mutant (*fbn2b^-/-^*) zebrafish had a reproducible phenotype, with a subset of embryos showing endocardial detachment leading to early mortality. Interestingly, the remaining *fbn2b*^-/-^ zebrafish developed dilation of the bulbus arteriosus, a structure analogous to the aortic root in humans, and survived normally to adulthood. Adult *fbn2b^-/-^* zebrafish displayed cardiac valve abnormalities. Transcriptomic analysis of *fbn2b^-/-^*embryos suggested the involvement of extracellular matrix remodeling and immune-related pathways. Administration of nebivolol and losartan did not improve the phenotype of *fbn2b^-/-^* larvae. Zebrafish lacking fibrillin-1 and/or fibrillin-2a did not show any phenotype.

**Conclusion:** Our *fbn2b^-/-^* zebrafish model recapitulates key aspects of human cardiovascular manifestations of MFS and can therefore be considered a novel relevant animal model for MFS. Studying this model allows us to broaden the knowledge of the underlying mechanisms of the disease and discover much-needed disease-specific treatment options.

**CONDENSED ABSTRACT:** Fibrillin defects lead to severe cardiovascular complications in Marfan syndrome (MFS), including aortic dilation, dissection, and rupture. To model MFS, we generated zebrafish mutants lacking various fibrillin genes. Among these mutant lines, only fibrillin-2b-deficient zebrafish exhibited cardiovascular phenotypes mimicking human disease. Multimodal imaging revealed early cardiac defects, bulbus arteriosus dilation, and valve abnormalities. Transcriptomic analysis identified altered regulation of pathways related to extracellular matrix homeostasis and immune system activation. Compound testing demonstrated the model’s potential for drug discovery. This zebrafish model, recapitulating key cardiovascular features of MFS, provides a valuable platform to investigate disease mechanisms and identify novel treatment strategies.

## INTRODUCTION

Fibrillin microfibrils are fundamental components of the extracellular matrix that contribute to the integrity of connective tissue in various organs, including blood vessels, lungs, skin, skeleton, and eyes.^1^ These microfibrils can either directly provide stress-bearing structural support to the tissue or play an essential role as a scaffold for tropoelastin deposition, leading to the formation of elastic fibers that provide tensile strength to the extracellular matrix.^2,3^ In addition to their structural role, fibrillin microfibrils are also crucial for tissue mechanobiology and homeostasis through the regulation of the bioavailability of growth factors of the transforming growth factor-β (TGF-β) and bone morphogenetic protein family, and by interactions with cell surface receptors such as integrins.^4^

The human genome contains three fibrillin isoforms: fibrillin-1 (*FBN1*), −2 (*FBN2*) and −3 (*FBN3*). Pathogenic variants in *FBN1* and *FBN2* have been associated with connective tissue disorders, including Marfan syndrome (MFS, OMIM #154700) and Beals-Hecht syndrome, also known as congenital contractural arachnodactyly (OMIM #121050), respectively.^5–8^ MFS is an autosomal dominant inherited disorder with pleiotropic manifestations, including skeletal abnormalities (e.g. skeletal overgrowth, joint laxity), ocular manifestations (e.g. ectopia lentis), and skin abnormalities (e.g. striae).^1^ In addition to the cardinal cardiovascular manifestations – namely thoracic aortic aneurysm and dissection (TAAD) and mitral valve disease – impaired myocardial function and arrhythmias occur more frequently in patients with MFS. These cardiovascular manifestations contribute to significant morbidity and an increased risk of early mortality in patients with MFS.^9,10^ Patients with congenital contractural arachnodactyly have a MFS-like skeletal phenotype, but the eyes and aorta are typically not affected.^11^ Nevertheless, some cases have been described where pathogenic variants in human *FBN2* were linked to aortic dilation, indicating a level of functional overlap between different fibrillins.^12–14^ Noteworthy, syndromes with opposite phenotypes (e.g. short instead of tall stature) have been observed as well in patients harbouring pathogenic variants in specific domains of *FBN1* and *FBN2*.^15^ To date, the importance of *FBN3* remains less well characterized. However, some associations of *FBN3* variants were reported with polycystic ovary syndrome, Bardet-Biedl syndrome, and Weill-Marchesani syndrome.^16,17^

Previous research on MFS and other fibrillinopathies has significantly advanced our understanding of their genetic basis and the importance of fibrillin proteins for maintaining extracellular matrix integrity and biomechanical signalling. Nevertheless, the precise molecular mechanisms linking these fibrillin defects to the complex cardiovascular manifestations remain incompletely understood. There is thus a particular need for more flexible *in vivo* models to address this knowledge gap.

During the last decades, zebrafish (*Danio rerio*) has emerged as a versatile animal model to complement established mammalian models. Although the cardiovascular system of teleosts has a less complex architecture than its mammalian counterpart, mainly due to the lack of pulmonary circulation, it has nonetheless proven to be a valuable model for studying cardiovascular diseases.^18,19^ Zebrafish have several unique characteristics which make them an attractive disease model: (1) optical transparency during early development, enabling easy intravital microscopic observation, (2) suitability for high-throughput applications due to low cost and high fecundity, and (3) high genetic similarity to humans with over 70% of human coding genes having at least one zebrafish orthologue.^20^ For cardiac pathologies specifically, 96% of the genes known to cause cardiomyopathies are conserved and highly expressed in the zebrafish heart.^21^ These distinctive traits make zebrafish an invaluable model for cardiovascular research, particularly for small-molecule high-throughput drug screens to discover novel therapeutic targets.^19^ Due to their genetic tractability, zebrafish can also be used to assess the physiological effects of variants of uncertain significance, which would be a significant contribution to improved patient management.^22,23^

Like humans, the zebrafish genome contains three fibrillin genes, termed *fibrillin-1* (*fbn1*), *-2a* (*fbn2a*), and *-2b* (*fbn2b*), though conflicting annotations have been used in different publications and databases. A phylogenetic study conducted by Piha-Gossack et al.^24^ has revealed only small evolutionary changes in fibrillin protein structure among different species. The ancestral fibrillin gene already contained most key elements except for the unique characteristic domain located at the N-terminus and specific RGD (Arg-Gly-Asp) motifs.

In this study, we systematically disrupted the three different fibrillin genes in zebrafish using CRISPR-Cas9 to examine their impact on the development and function of the cardiovascular system. Our findings revealed multiple phenotypes that are pertinent to MFS.

## MATERIAL AND METHODS

All materials and methods are described in detail in the Supplemental Methods.

## RESULTS

### Protein homology between human and zebrafish fibrillin isoforms

The protein structure of the human fibrillin family has been well-documented and shows that the different fibrillin genes share an almost identical organization of functional domains. To analyze the zebrafish fibrillin gene family, we used the genomic *fbn2a* and *fbn2b* sequences, which were mapped and annotated in the GRCz11 assembly of the zebrafish genome sequence. A genome assembly gap, however, prevented correct annotation of the first 32 exons of the zebrafish *fbn1* gene. Recently, a full-length cDNA sequence prediction was assigned to *fbn1* (XM_073930076.1), and this sequence, which aligned with our own cDNA sequencing data, was used for in-depth analysis (Supplemental Figure 1). The zebrafish fibrillins present a high level of homology to human fibrillin proteins with an average overlap in amino acid sequence of 70% (Figure 1B). Further supporting this, InterProScan analysis revealed a highly conserved domain organization between species, with the characteristic 4-cysteine domain, two hybrid domains, and seven 8-cystein or TGF-β binding-like domains present in all identified zebrafish fibrillin proteins (Figure 1A).

**Figure 1.**
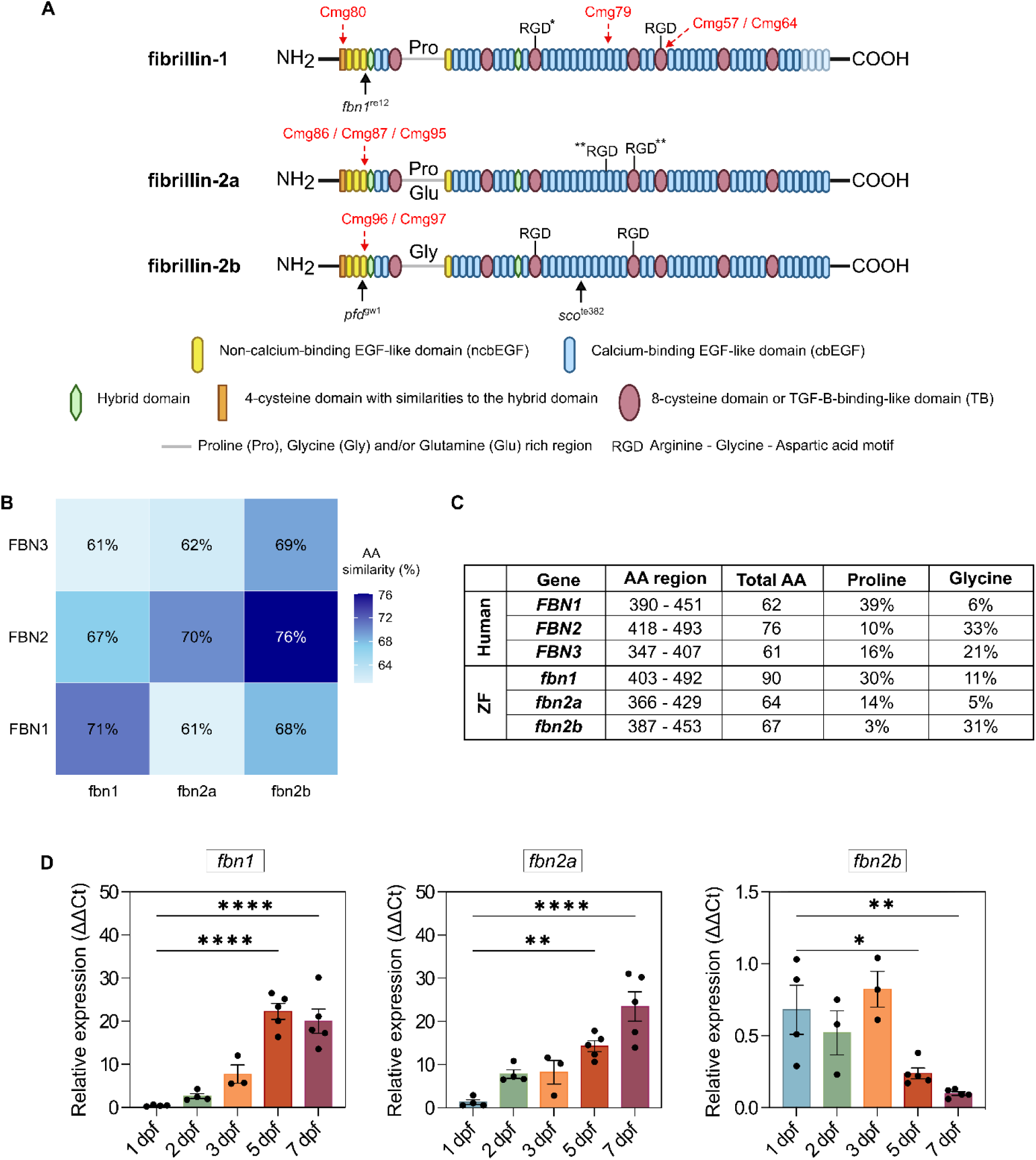
Overview of the three fibrillin isoforms in zebrafish. **(A)** Schematic representation of the protein domains identified in all three zebrafish fibrillin isoforms (fibrillin-1, 2a, and 2b). Unique Pro/Gly/Glu-rich regions and the various RGD-motifs are indicated, of which some are exclusively present in zebrafish (*). CRISPR/Cas9-induced recombination sites are indicated with a red arrow; mutation sites in previously reported zebrafish models are also annotated with a black arrow. (B) Heatmap showing the percentage of amino acid sequence similarity between human fibrillin proteins (Y-axis) and their zebrafish orthologs (X-axis). Sequence similarity was calculated using Clustal Omega. **(C)** Comparison of the relative contribution of proline (Pro) and glycine (Gly) in the Pro/Gly-rich region between human and zebrafish fibrillins. **(D)** mRNA expression pattern of the fibrillin isoforms in WT zebrafish embryos at 1, 2, 3, 5 and 7 dpf (n = 3-5 for each developmental stage). Data are expressed as a mean ± SEM. Statistical test analysis: one-way ANOVA followed by Dunnett’s multiple comparisons test on log-transformed data. ****p < 0.0001, ***p < 0.001 and **p < 0.01. AA = amino acid, ZF = zebrafish.

A characteristic domain within all fibrillins is the proline and/or glycine-rich region immediately following the first TGF-β binding-like domain. Comparing these domains between species reveals similarities in amino acid composition between *FBN1* and *fbn1*, *FBN2* and *fbn2b*, and, to some extent, *FBN3* and *fbn2a* (Figure 1C). Furthermore, the RGD-integrin binding sites present in all human fibrillin isoforms are highly conserved in the zebrafish fibrillins, except for zebrafish fibrillin-2a, which lacks all RGD motifs. Interestingly, fibrillin-1 in zebrafish contains an extra RGD motif in the third TGF-β binding-like domain. The protein structures of all three zebrafish fibrillins are summarized in Figure 1A.

### Expression pattern of the different fibrillin isoforms in wild-type (WT) zebrafish

We used real-time qPCR to measure expression levels of *fbn1*, *fbn2a* and *fbn2b* in WT whole embryo tissue at different developmental stages (1, 2, 3, 5 and 7 dpf). Our results showed that both *fbn1* and *fbn2a* are expressed more abundantly later in development, whereas *fbn2b* expression levels are highest at the early stages and gradually decrease with age (Figure 1D).

### Normal cardiovascular development in zebrafish lacking fbn1 and/or fbn2a

Four independent *fbn1* knock-out (KO) zebrafish models were generated, carrying either a frameshift deletion or insertion in exon 2, 34, or 38, as summarized in Supplemental Table 1. Surprisingly, all *fbn1* homozygous mutant (*fbn1^-/-^*) zebrafish survived normally to adulthood, without any cardiovascular phenotype during development (Figure 2A, 2D). Loss of *fbn2a*, with or without *fbn1* deficiency, also did not influence cardiovascular development or survival (Figure 2A). Real-time qPCR expression analysis demonstrated a significant reduction in *fbn1* mRNA expression in *fbn1^-/-^* zebrafish compared to WT siblings, observed in both 5 dpf whole larvae (Figure 2B) and in adult (9 mpf) ocular, skin, and muscle tissues (Figure 2C). *fbn2a* and *fbn2b* mRNA expression was not affected in *fbn1* mutants (Supplemental Figure 2). Transthoracic echocardiography performed on adult *fbn1* and/or *fbn2a* mutant zebrafish of various ages (6 – 19 mpf) also demonstrated no significant abnormalities in any measured cardiovascular parameters (Supplemental Tables 6 and 7).

**Figure 2.**
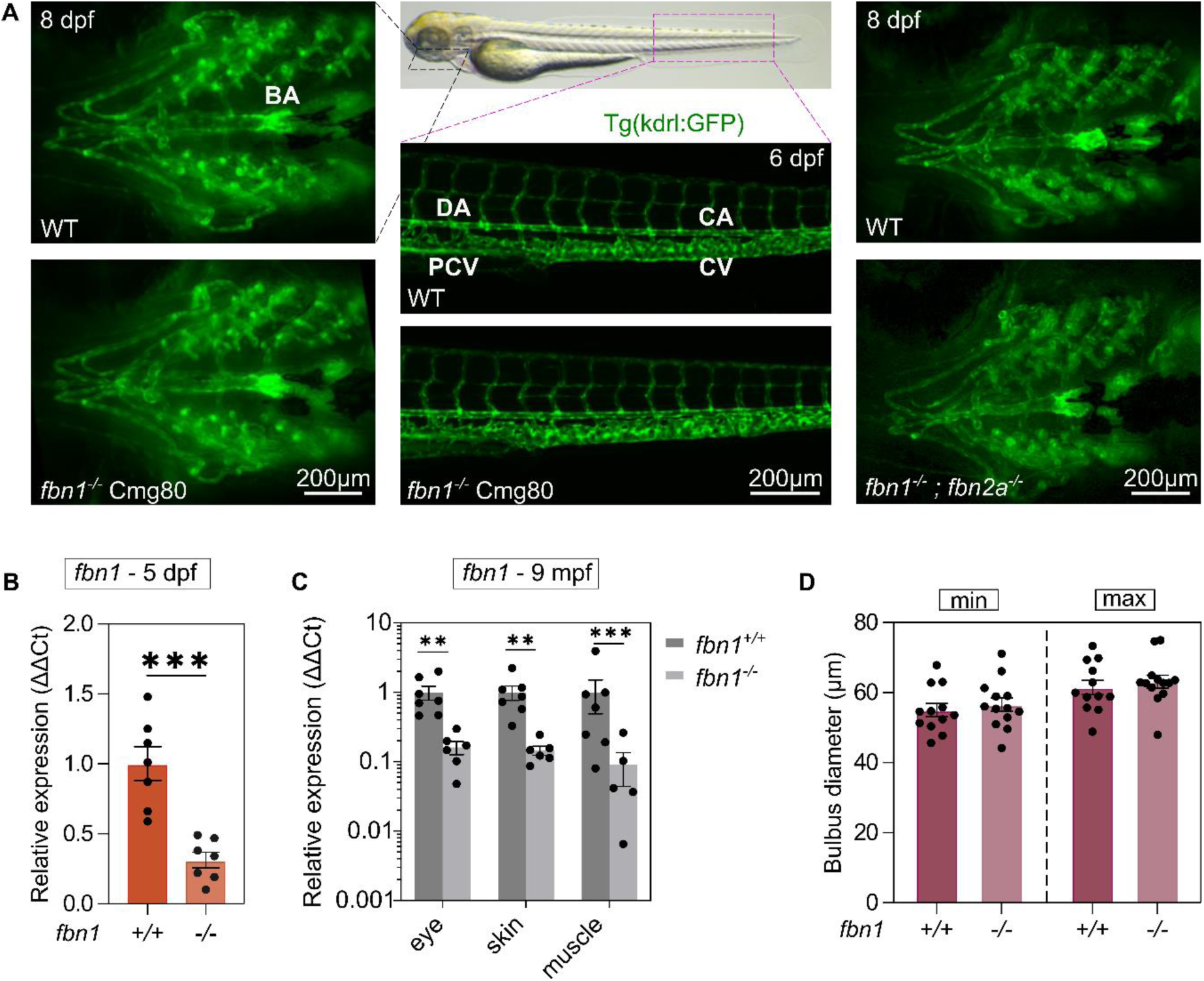
Cardiovascular architecture in fbn1 and/or fbn2a mutants. Fluorescent images of the vasculature of *fbn1^-/-^* (Cmg80) zebrafish with or without the additional loss of *fbn2a* at 8 dpf, showed no phenotypic differences from WT. **(A – left and right)** Ventral view of the ventral aorta (VA) and BA of 6-8 dpf WT, *fbn1^-/-^*and *fbn1^-/-^;fbn2a^-/-^* larvae respectively. **(A - middle)** Lateral view of the distal part of the dorsal aorta (DA) merging into the caudal aorta (CA) as well as the posterior cardinal vein (PCV) merging into caudal vein (CV) in 6 dpf WT and *fbn1^-/-^* larvae. (**B)** RT-qPCR analysis of *fbn1* expression in 5 dpf WT and *fbn1^-/-^*(Cmg80) larvae (n = 7). Each data point represents the mean of two technical repeats. (**C)** RT-qPCR analysis of *fbn1* expression in eye, skin, and muscle tissue of 9 mpf WT and *fbn1^-/-^* (Cmg80) zebrafish (n = 5-7). **(D)** Quantification of BA diameters in 7 dpf *fbn1^-/-^* (Cmg80) and matched WT controls during minimal (min) and maximal (max) distension (n = 12 – 13). Data are expressed as mean ± SEM. Statistical analysis: unpaired t-test (B), two-way ANOVA (C, D). ***p<0.001, ***p<0.01.

### Variable endocardial phenotype in fbn2b^-/-^ zebrafish

Next, we generated a 4 bp frameshift deletion located in exon 4 of the *fbn2b* gene, resulting in a premature termination codon (p.Cys153*). Real-time qPCR expression analysis of *fbn2b^-/-^* larvae (1, 2, 3, 5 and 7 dpf) showed a strong and significant reduction of *fbn2b* mRNA expression compared to WT siblings, likely due to nonsense-mediated decay (Figure 3E + Supplemental Figure 3). *fbn1* and *fbn2a* mRNA expression was not affected in *fbn2b* mutants (Supplemental Figure 3). Starting at 24 hpf, *fbn2b* homozygous mutant (*fbn2b^-/-^*) zebrafish embryos can be distinguished from their heterozygous and WT siblings by the presence of fin fold atrophy. In approximately 40% of *fbn2b^-/-^* offspring, marked pericardial edema was observed, associated with gaps in the endocardium which progress to complete endocardial detachment in the atrium, similar to what was previously reported for the *scotch tape* ENU *fbn2b* mutant (Figure 3A).^25^ *fbn2b^-/-^* zebrafish with this severe phenotype do not survive past 6-8 dpf due to vascular embolism. The remaining *fbn2b^-/-^* larvae exhibit a milder phenotype where the endocardium remains attached to the myocardium, circulation is maintained, and normal survival to adulthood is observed (Figure 3A). Interestingly, we observed that a subset of *fbn2b^-/-^*embryos with severe pericardial edema (approximately 30%) seems to recover between 3-5 dpf, reverting to the milder phenotype (Figure 3B).

**Figure 3.**
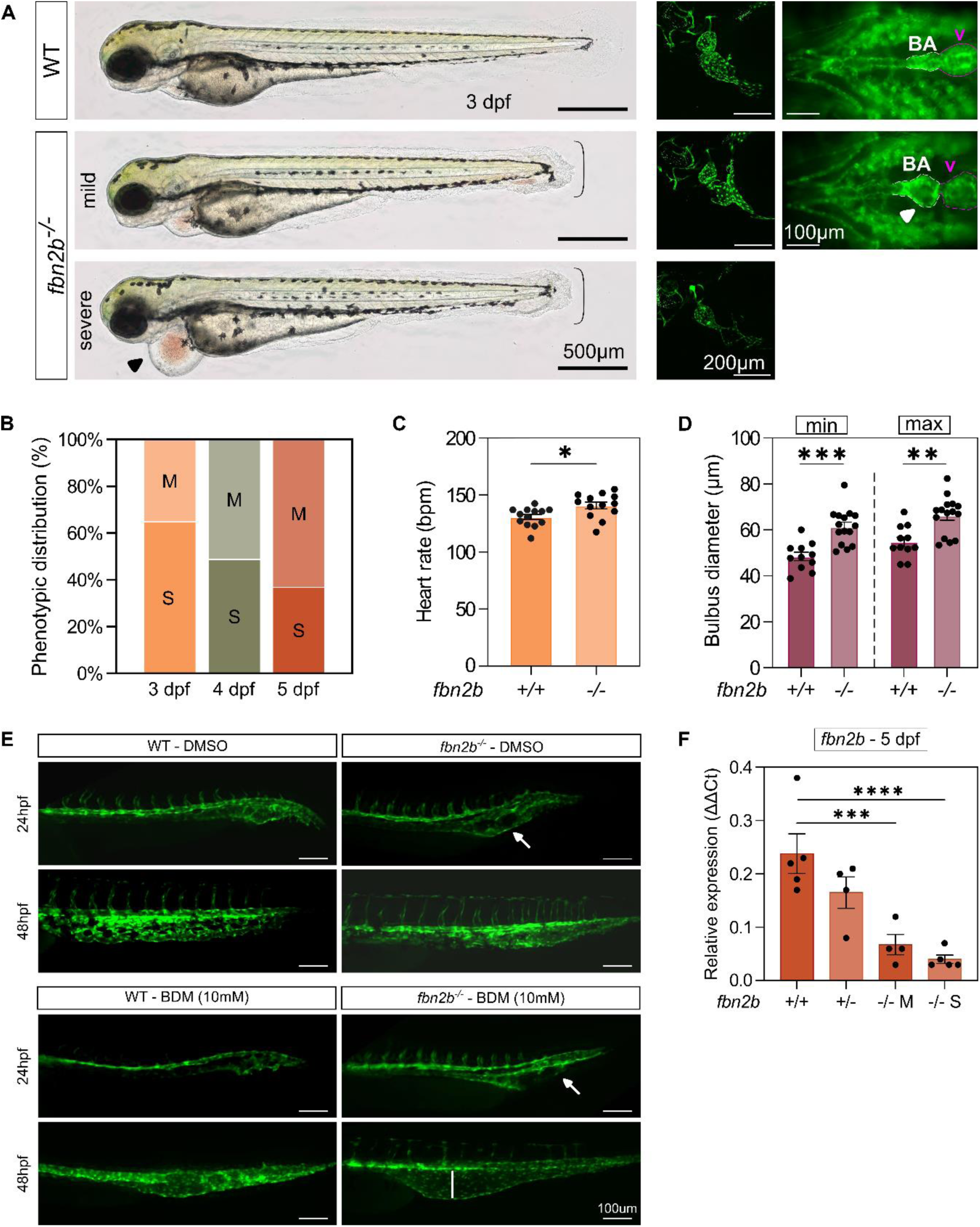
Cardiovascular development in fbn2b mutants. Representative images of the diverse phenotypes observed in 2-8 dpf *fbn2b^-/-^* (cmg96) mutants and WT controls. **(A - left)** Lateral whole-embryo view of 3 dpf larvae using brightfield microscopy. Accolade indicates finfold atrophy, black arrowhead indicates severe pericardial edema. **(A - middle)** Reconstructed 3D *in vivo* two-photon fluorescent images of the non-beating heart of 2 dpf *Tg(kdrl:GFP)* WT and *fbn2b^-/-^* zebrafish. Endocardial detachment (asterisk) is observed in the atrium of the *fbn2b^-/-^* zebrafish with pericardial edema. **(A - right)** Ventral view of 8 dpf *Tg(kdrl:GFP)* WT and *fbn2b^-/-^* with preserved endocardial integrity. White arrowhead indicates dilated BA. **(B)** Phenotypic distribution (%) of *fbn2b*-deficient zebrafish presenting a mild (M) or severe (S) pericardial phenotype and their dynamics over time (3, 4 and 5 dpf) (n = average of 19 clutches). **(C)** Quantification of the average heart rate in beats per min (bpm) of 3 dpf *fbn2b^-/-^*with preserved endocardial integrity and matching controls (n = 13). **(D)** Quantification of BA diameters at 7 dpf during minimal (min) and maximal (max) distension (n = 11-15). **(E)** Caudal vein formation in *Tg(kdrl:GFP)* WT and mild *fbn2b^-/-^* zebrafish at 24 and 48 hpf after exposure to 0.01% DMSO vehicle (top) or 10 mM 2,3-butanedione monoxime (BDM) to inhibit cardiac contraction (bottom). White arrow indicates abnormal development of the caudal vein. Exposure to BDM leads to more severe caudal vein dilatation (white line) in *fbn2b^-/-^* embryos than in WT controls at 48 hpf (qualitative analysis). **(F)** mRNA expression levels of *fbn2b* in WT, *fbn2b^+/-^*, and *fbn2b^-/-^* (mild or severe) zebrafish at 5 dpf (n = 4-5). Each datapoint represents the mean of two technical replicates. Statistical analysis: unpaired t-test (C), two-way ANOVA (D), one-way ANOVA followed by Tukey multiple comparison’s test on log-transformed data. Data are expressed as mean ± SEM. ****p<0.0001, ***p<0.001, **p<0.01, *p<0.05. BA = bulbus arteriosus, v = ventricle.

### Bulbus arteriosus and cardiac function in fbn2b^-/-^ zebrafish larvae

Interestingly, the surviving *fbn2b^-/-^* zebrafish in which the endocardium remains attached develop dilatation of the bulbus arteriosus (BA), a structure that is considered to be evolutionarily related to the aortic root and ascending aorta in humans, starting at 5 dpf (Figure 3A, 3D). We also tested several cardiac function parameters using brightfield microscopy and found a mild but significant increase in heart rate in mild *fbn2b^-/-^* (without pericardial edema) compared to WT controls at 3 dpf (Figure 3C, Supplemental Figure 4).

### Venous phenotype in fbn2b^-/-^ zebrafish larvae

Besides the BA phenotype, we also discovered that the caudal vein of *fbn2b^-/-^* zebrafish initially develops as a dilated, cavernous venous structure lacking vessel integrity. While this phenotype persists in the *fbn2b* mutants with the severe endocardial phenotype, we found that, in the mild *fbn2b* mutants, the caudal vein remodels appropriately (Figure 3E). This led us to investigate the role of blood flow dynamics further by pharmacologically inhibiting cardiac contraction using a myosin inhibitor. This prevented the resolution of the pronounced caudal vein dilatation in the mild *fbn2b^-/-^* zebrafish (Figure 3E).

### qPCR confirms the absence of compensation mechanisms by fbn1 and/or fbn2a

To test whether other fibrillin isoforms compensate for a lack of *fbn2b* expression in the *fbn2b^-/-^* zebrafish, the expression levels of *fbn1*, *fbn2a* and *fbn2b* were measured in WT, *fbn2b^+/-^*, *fbn2b^-/-^* mild and *fbn2b^-/-^* severe whole embryos during early development (1, 2, 3, 5 and 7 dpf). No significant differences in *fbn2b* expression levels were observed between the mild and severe *fbn2b^-/-^* phenotypes (Figure 3F, Supplemental Figure 3). Also, no differences were observed in *fbn1* and *fbn2a* expression levels between *fbn2b^-/-^* and matching controls, nor between *fbn2b^-/-^* larvae with a mild versus severe phenotype (Supplemental Figure 3). This suggests that increased expression of *fbn1* or *fbn2a* does not compensate for the lack of *fbn2b* expression.

### Triple fibrillin knockouts do not survive to adulthood

Since loss of *fbn1* and/or *fbn2a* does not lead to a phenotype, we investigated whether additional loss of *fbn2b* could unmask a functional role for these fibrillins. Consistent with the single *fbn2b* KO phenotype, triple knock-out (TKO) mutants exhibited 100% penetrance of finfold atrophy, and a subset of TKO larvae demonstrated severe pericardial edema due to complete endocardial detachment while the remaining TKO had a milder phenotype. (Figure 4A, 4B). Kaplan-Meier survival analysis of offspring from a *fbn1^-/-^;fbn2a^-/-^;fbn2b^+/-^* incross up to 14 dpf showed slightly lower survival rates in TKO in comparison to siblings, although statistical significance was not reached (Figure 4C). To date, we have, however, never detected an adult TKO zebrafish despite genotyping approximately 300-400 adult fish raised from *fbn1^-/-^;fbn2a^-/-^;fbn2b^+/-^* incrosses, indicating premature mortality of zebrafish with this genotype. We next assessed the standard length (SL) at 3 dpf and found that TKO mutants are significantly shorter than their *fbn1^-/-^;fbn2a^-/-^;fbn2b^+/+^* siblings (Figure 4E). Finally, measurements of the BA diameter at 7 dpf revealed a significant increase in the minimally distended diameter in TKO larvae compared to *fbn1^-/-^;fbn2a^-/-^;fbn2b^+/+^* controls. In the maximally distended state, a larger variability was observed in the TKO group (Figure 4D).

**Figure 4.**
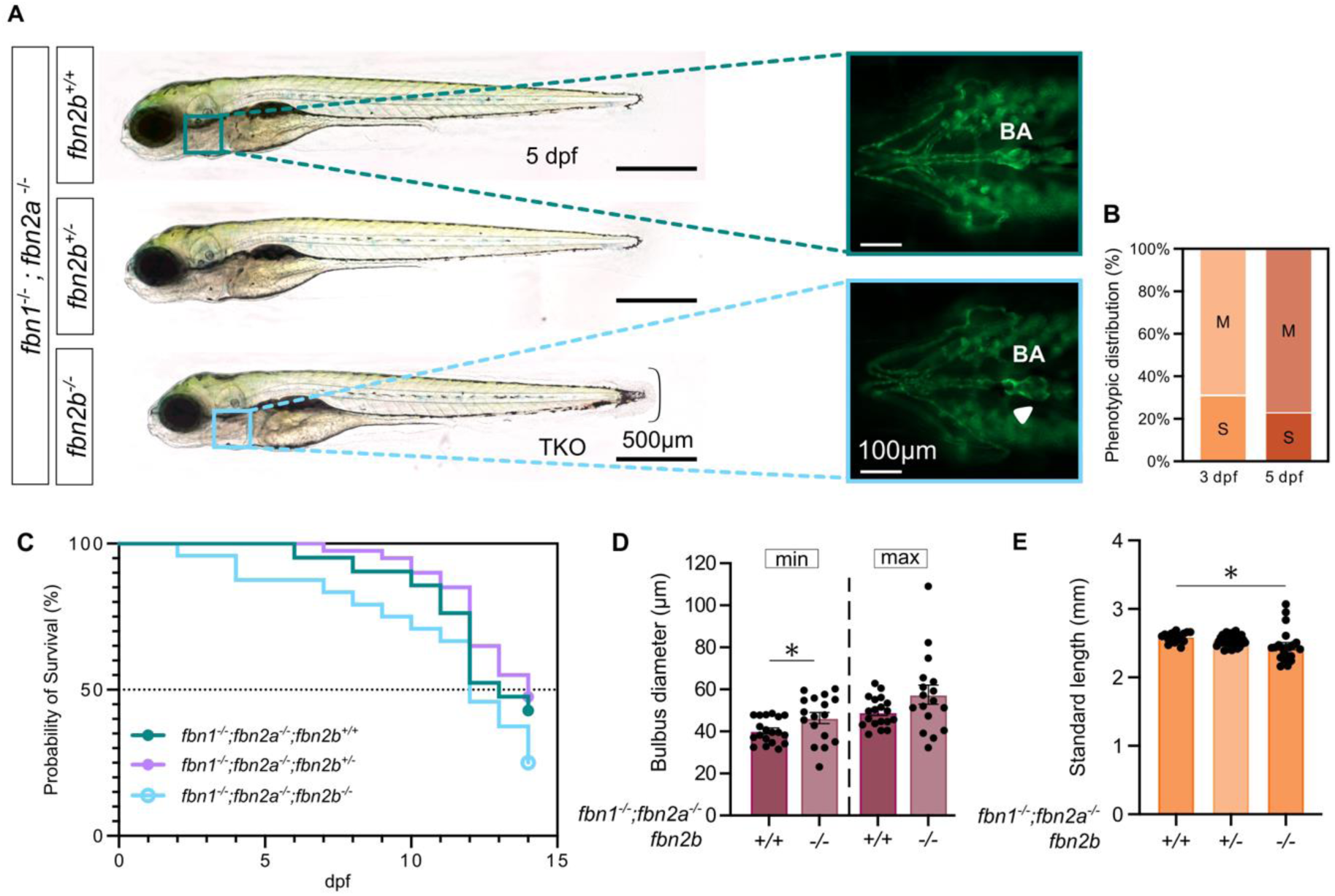
Phenotypic features of TKO larvae. Representative images of the phenotypes observed in 3-7 dpf *fbn1^-/-^;fbn2a^-/-^;fbn2b^+/+^*, *fbn1^-/-^;fbn2a^-/-^;fbn2b^+/-^, and fbn1^-/-^;fbn2a^-/-^;fbn2b^-/-^*larvae. **(A – left)** Lateral whole-embryo view of a 5 dpf triple fibrillin knockout (TKO) without complete endocardial detachment and sibling controls, using brightfield microscopy. Accolade: finfold atrophy. **(A – right)** Fluorescent ventral images of 7 dpf *Tg(kdrl:GFP)* TKO and sibling controls with preserved endocardial integrity. White arrowhead indicates dilated BA. **(B)** Phenotypic distribution (%) of *TKO* zebrafish presenting a mild (M) or severe (S) pericardial phenotype and their dynamics over time (3 and 5 dpf) (n = 21). **(C)** 14-day Kaplan-Meier survival curve of the offspring of an incross of *fbn1^-/-^;fbn2a^-/-^;fbn2b^+/-^* zebrafish (n = 21-40). **(C)** Quantification of BA diameters at 7 dpf during minimal (min) and maximal (max) distension (n = 17-19). **(D)** Quantification of standard length at 3 dpf (n = 17 – 36). Statistical analysis: one-way Anova followed by Dunnett’s multiple comparison’s test. All data are expressed as a mean ± SEM. *p<0.05. BA = bulbus arteriosus, dpf = days post fertilization.

### Transcriptomic analysis in fbn2b^-/-^ embryos suggests involvement of extracellular matrix remodeling and immune system activation

We conducted bulk RNA sequencing on whole *fbn2b^-/-^* and WT embryos at 1 and 2 dpf, to identify early transcriptional changes during the development of the early cardiovascular phenotypes. Differential expression analysis (adjusted p-value (FDR) < 0.05 and log_2_FC > 1) identified 42 and 361 differentially expressed genes, at 1 and 2 dpf, respectively (Figure 5A, 5C, Supplemental Table 8). At 1 dpf, gene ontology enrichment analysis of the differentially expressed genes revealed significant enrichment in pathways related to metabolism, biosynthesis, and endothelial and cardiac tissue development (FDR < 0.05) (Figure 5B, 5D). The latter transcriptional changes were notably driven by strong downregulation of the *fbn2b* transcript (log_2_FC-3.22). By 2 dpf, the transcriptional profile shifted, with gene ontology analysis highlighting enrichment of immune-related processes and defence mechanisms. In particular, several components of the complement system were significantly upregulated in *fbn2b^-/-^* mutants, including *c4b* (log_2_FC +1.37), *c6* (log_2_FC +3.69), *cfb* (log_2_FC +1.11), *c7a* (log_2_FC +0.94), and *c7b* (log_2_FC +3.18). Additionally, matrix metalloproteinases were markedly increased, with elevated expression of *mmp9* (log_2_FC +2.77), *mmp13a* (log_2_FC +2.57) and *mmp13b* (log_2_FC +2.85) (Figure 5).

**Figure 5.**
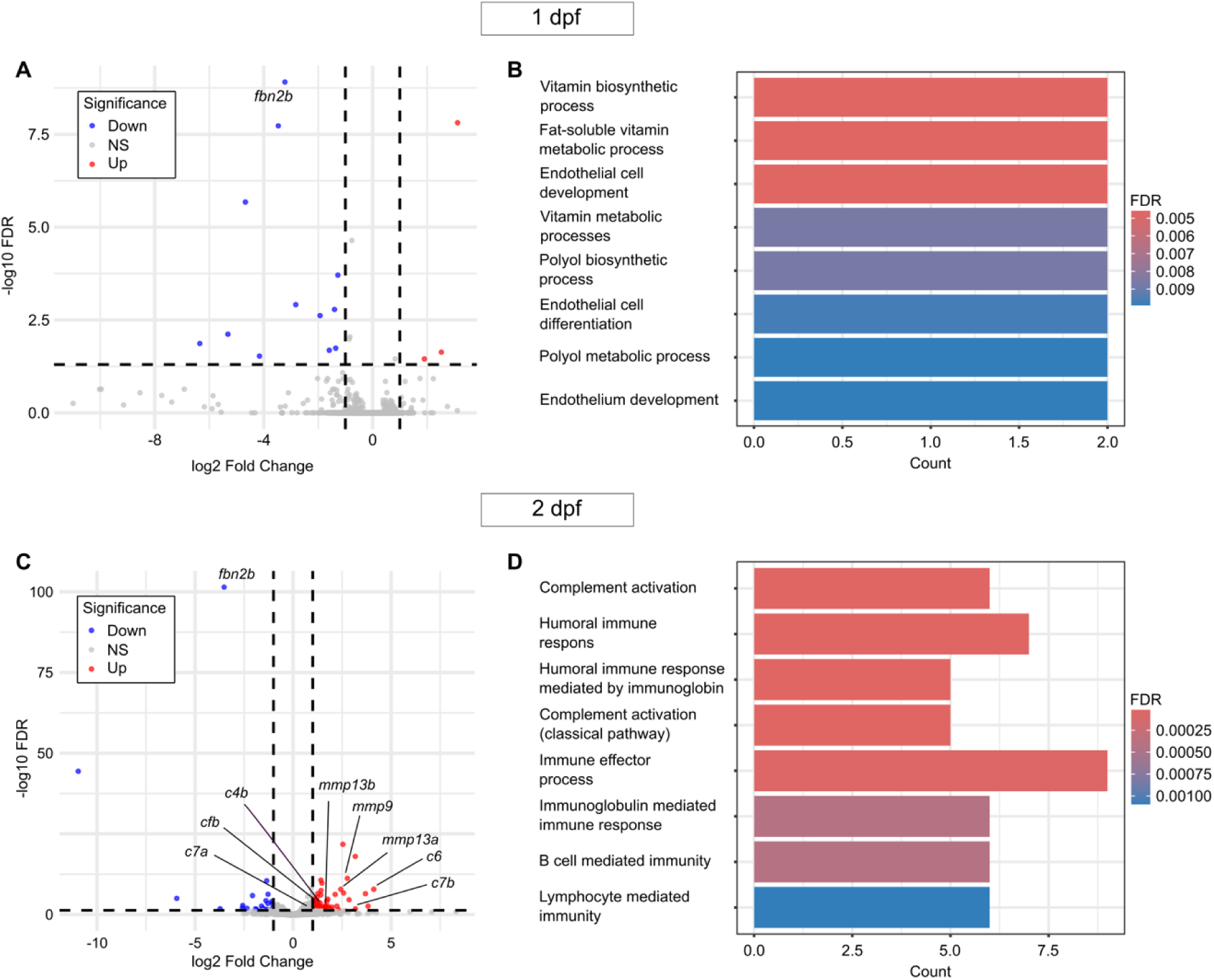
Transcriptomic analysis of fbn2b^+/+^ and fbn2b^-/-^ siblings at 1 and 2 dpf. **(A, C)** Volcano plots illustrating differentially expressed genes (DEG) in *fbn2b^-/-^* zebrafish compared to WT siblings at 1 dpf and 2 dpf, respectively. Upregulated genes are red and downregulated genes are blue, with genes of importance annotated. Thresholds: FDR ≥ 0.05 and |log2FC| ≥ 1. **(C, D)** GO enrichment analysis with top 8 hits presented as a barplot. The x-axis and y-axis represent the gene count and pathway, respectively. Thresholds: FDR ≥ 0.05 and |log2FC| ≥ 1.

### Echocardiographic and synchrotron imaging in adult fbn2b^-/-^ zebrafish hearts reveal cardiac rhythm and morphological abnormalities

Using an optimised in-house cardiac ultrasound setup, we assessed the cardiovascular phenotype of adult *fbn2b^-/-^* zebrafish *in vivo*. We detected significant dilation of BA of 6 and 8-month-old *fbn2b* mutants compared to WT zebrafish (Figure 6B, 6D). Additionally, the ventricles of the *fbn2b* mutants exhibited mild dilation (Figure 6B, 6C). Analysis of color flow Doppler (CFD) recordings revealed increased blood inflow and outflow areas in *fbn2b^-/-^* zebrafish, indicating that AV and BV valve openings are larger than in WT zebrafish. A small number of *fbn2b^-/-^* zebrafish showed valvular regurgitation (Figure 6F, 6G).

**Figure 6.**
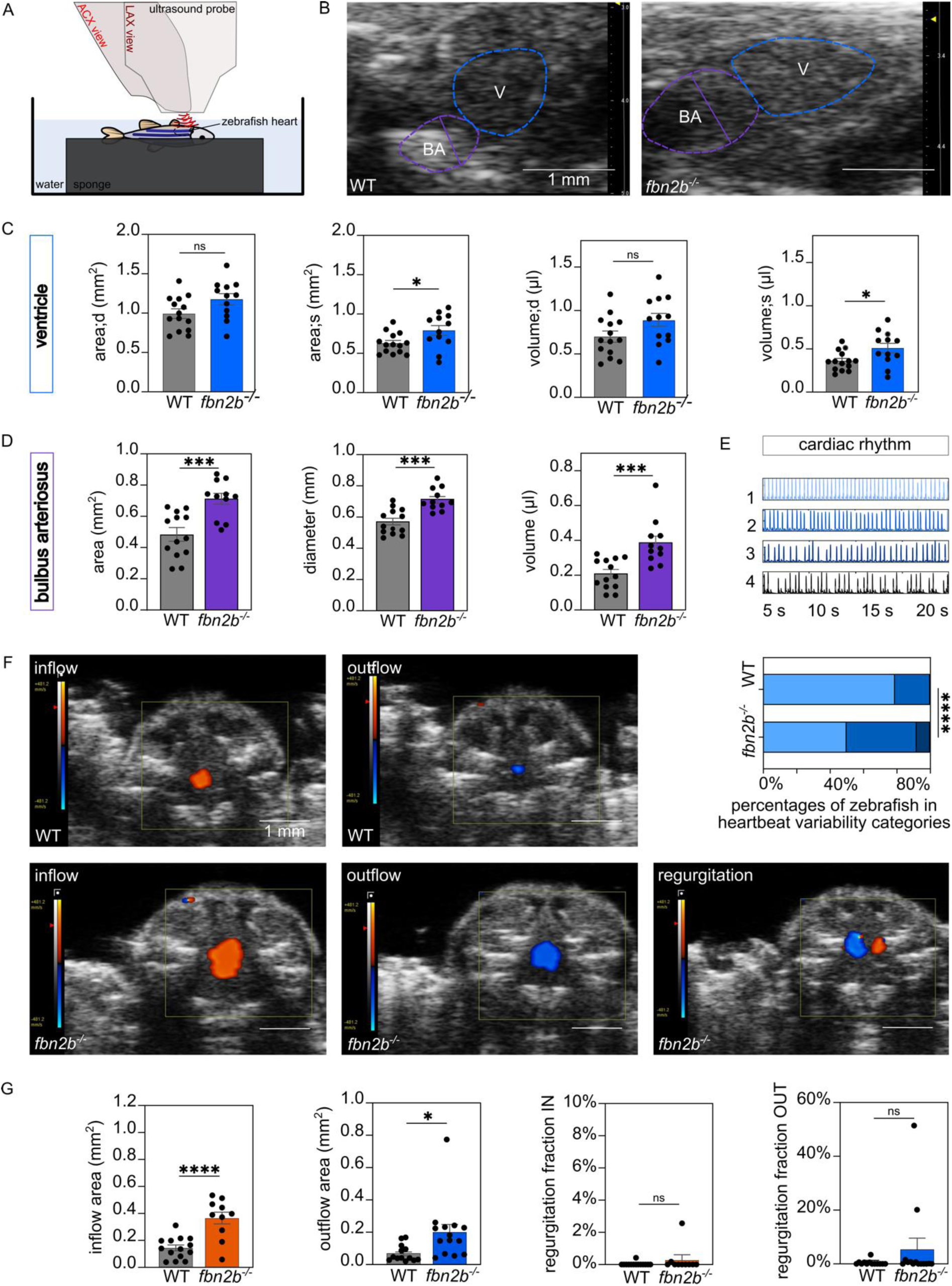
Cardiac abnormalities of adult fbn2b^-/-^ zebrafish. **(A)** Schematic representation of the two-dimensional transthoracic echocardiography of adult zebrafish. The zebrafish is anesthetized, positioned with its ventral side upwards, and submerged in water. The echocardiography measurements are made with an ultrasound probe, in abdominocranial axis view for CFD analysis (red), and in longitudinal view for ventricle and BA measurements (dark red). **(B)** Tracings of the posterior walls of the ventricle (blue) and BA (purple) of WT (left) and *fbn2b^-/-^* zebrafish (right) (6 and 8 mpf). **(C)** Dimensions of the ventricle during diastole (area;d, volume;d) and systole (area;s, volume;s). **(D)** Measurements of BA volume, area, and diameter while in maximal relaxation (at the time of BV valve contraction). **(E)** Representative 20-second cardiac rhythm tracings with the associated qualitative score (top). Distribution of cardiac rhythm scores for different genotypes (n = 12) (bottom). **(F)** CFD recordings of inflow (orange) and outflow (blue) of blood into/from the ventricle in WT (top) and *fbn2b^-/-^* zebrafish (bottom), with an example of regurgitant blood flow seen in some mutants (2/14). **(G)** Quantification of inflow (orange) and outflow (blue) areas and regurgitation fractions from the CFD data. Data are expressed as mean ± SEM. Statistical test analysis: unpaired t-test (C, D, G – inflow area), Mann-Whitney test (G – outflow area, regurgitation fraction IN, regurgitation fraction OUT), Fischer’s exact test (E). ****p<0.0001, ***p<0.001, **p<0.01, *p<0.05, ns = non-significant. ACX = abdominocranial view, BA = bulbus arteriosus, LAX = longitudinal axis view, V = ventricle, WT = wild type.

We scored *fbn2b^-/-^* and WT zebrafish cardiac rhythm based on 20-second-long echocardiography recordings. *fbn2b^-/-^* zebrafish showed cardiac rhythm disorders, observed as more irregular or skipped heartbeats (Figure 6E).

By automated processing of PWD measurements, we observed no significant variations in cardiovascular parameters of systolic, diastolic, and valve function among *fbn2b* mutants compared to WT controls (Supplemental Table 6). When comparing the volumes of the atrium, ventricle, or BA between *fbn2b^-/-^* and WT zebrafish based on synchrotron imaging, we found no statistically significant differences. This *ex vivo* method couldn’t recapitulate the BA dilation, seen on echocardiography performed *in vivo*, likely due to a lack of intraluminal pressure postmortem. However, while comparing the 3D heart models of mutants and WT, we noticed increased variability in atrial volumes in *fbn2b^-/-^* hearts compared to WT hearts (F test: P = 0.011) (hearts - Supplemental Figure 6; BA – Supplemental Figure 9).

### Abnormal cardiac valve architecture in adult fbn2b^-/-^ zebrafish

Through histological staining for elastin of heart sections from 6 and 8-month-old *fbn2b* mutant zebrafish, we found a cardiac valve phenotype that is common in all *fbn2b* mutants. Specifically, all *fbn2b^-/-^* zebrafish display abnormalities in the bulboventricular (BV) valve, with one or both leaflets either more thickened or asymmetrically shaped, compared to the valves of WT zebrafish (Figure 7A). An altered morphology was also found in the atrioventricular (AV) valves of a small subset of *fbn2b^-/-^* zebrafish (2/12 samples), which showed excessive hypertrophy of the valve interstitial cells, leading to increased leaflet length and thickness.

**Figure 7.**
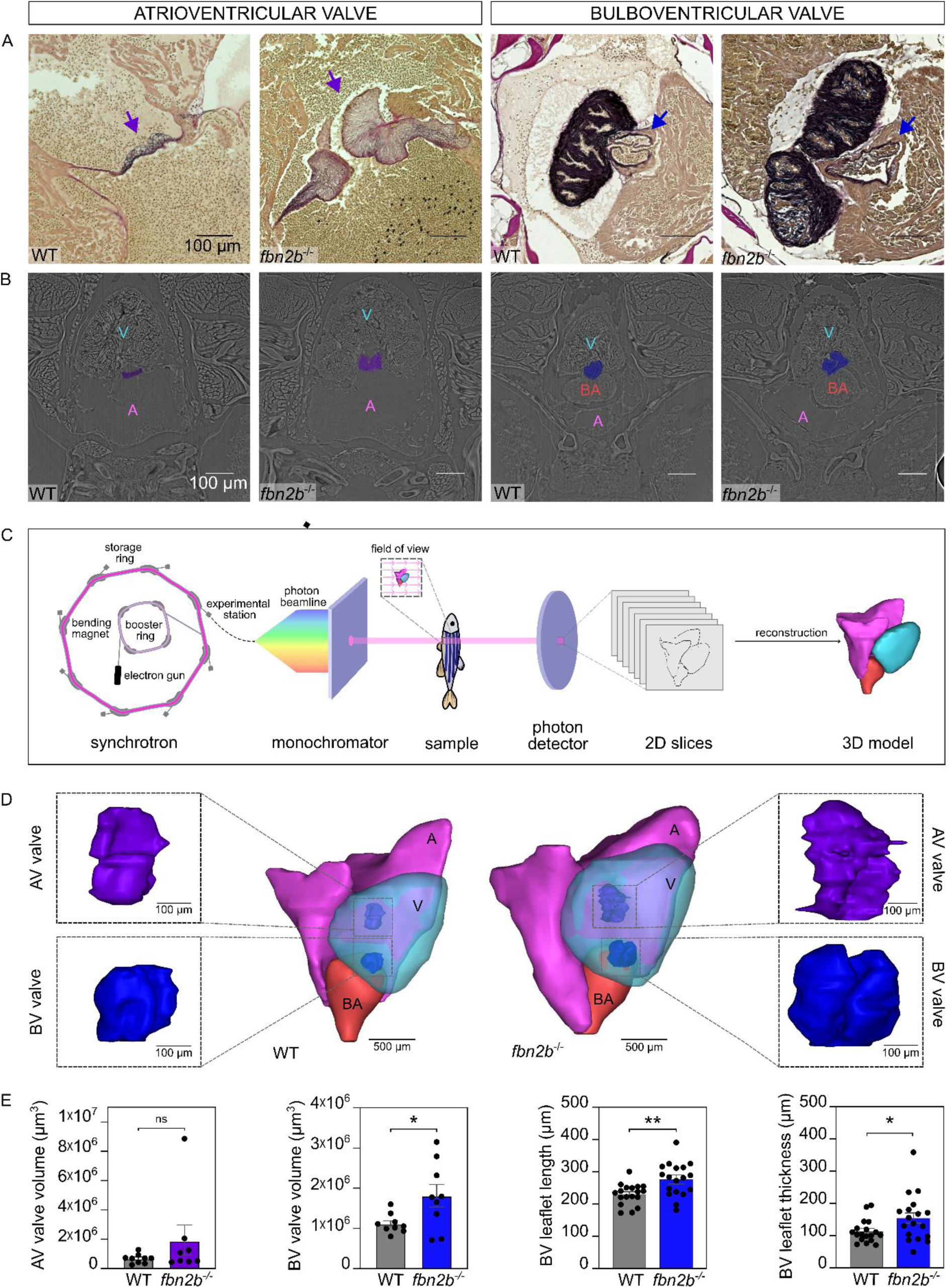
Abnormal cardiac valve architecture of adult fbn2b^-/-^ zebrafish. **(A)** Histological staining for elastin (purple) of AV (left, purple arrow) and BV valves (right, blue arrow) of WT and *fbn2b^-/-^* zebrafish (6 and 8 mpf). Extensive hypertrophy of the AV valve as shown in the representative *fbn2b^-/-^* image is found in a subset of mutants (2/12), while the BV valve leaflets are asymmetrical and abnormal in shape in all mutants. **(B)** Representative images of zebrafish hearts obtained by synchrotron X-ray scanning. Cardiac valves are shown in colour (AV valve – purple, BV valve – blue). **(C)** Schematic representation of synchrotron X-ray imaging method of zebrafish samples. **(D)** Representative 3D models of WT (left) and *fbn2b^-/-^* zebrafish (right) (16 and 18 mpf). Enlarged models of cardiac valves are shown to display the differences between WT and mutants. **(E)** Volume, length, and thickness measurements of AV (purple) and BV (blue) valves, obtained from the 3D models. Data are expressed as a mean ± SEM. **p<0.01, *p<0.05, ns = non-significant. Statistical test analysis: Mann-Whitney (AV valve volume), unpaired t-test (BV valve volume, BV leaflet length, BV leaflet thickness). A = atrium, AV = atrioventricular valve, BA = bulbus arteriosus, BV = bulboventricular valve, V = ventricle, WT = wild type.

Next, we studied the valves’ 3D structure through synchrotron X-ray imaging of hearts from *fbn2b^-/-^* and WT zebrafish (Figure 7D, AV valves – Supplemental Figure 7; BV valves – Supplemental Figure 8). We found that *fbn2b^-/-^* zebrafish have larger BV valve volumes, with the same tendency seen for AV valves. Indeed, the morphology of the AV valves in a subset of *fbn2b^-/-^* was extremely aberrant, with abnormal folding and fusing of various leaflet segments. Measurement of the length and thickness of individual BV leaflets confirmed abnormalities seen on histology, with leaflets of *fbn2b^-/-^* being longer and thicker than WT (Figure 7E).

### Similar response to β**-**adrenergic receptor modulation in early development of WT and fbn2b^-/-^ larvae

We tested β-adrenergic receptor modulation in WT and *fbn2b^-/-^* zebrafish (with a mild phenotype) in the first days of development. Epinephrine similarly elevated heart rate in WT and *fbn2b^-/-^* zebrafish, although the increase did not reach statistical significance in *fbn2b^-/-^*, likely due to the increased variability in this group (Supplemental Figure 10, top). Administration of the β-adrenergic receptor antagonist nebivolol consistently lowered heart rate in both WT and *fbn2b^-/-^* zebrafish during early development. In contrast, atenolol showed no significant impact on heart rate in either genotype (Supplemental Figure 10, bottom).

### Drug treatment with losartan and nebivolol does not improve the phenotype in fbn2b^-/-^ larvae

To verify the effects of β-adrenergic receptor and angiotensin II receptor blockade on the phenotype of our *fbn2b^-/-^* zebrafish model, we administered nebivolol at 1 µM and tested varying concentrations of losartan (100 – 1000 µM). There was no statistically significant improvement in the phenotypic distribution of any compound-treated *fbn2b^-/-^* zebrafish group at 3, 4, and 5 dpf compared to solvent-treated controls (Figure 8A). Similarly, no improvement of the dilated BA phenotype was observed in any group after 7 days of drug treatment (Figure 8B).

**Figure 8.**
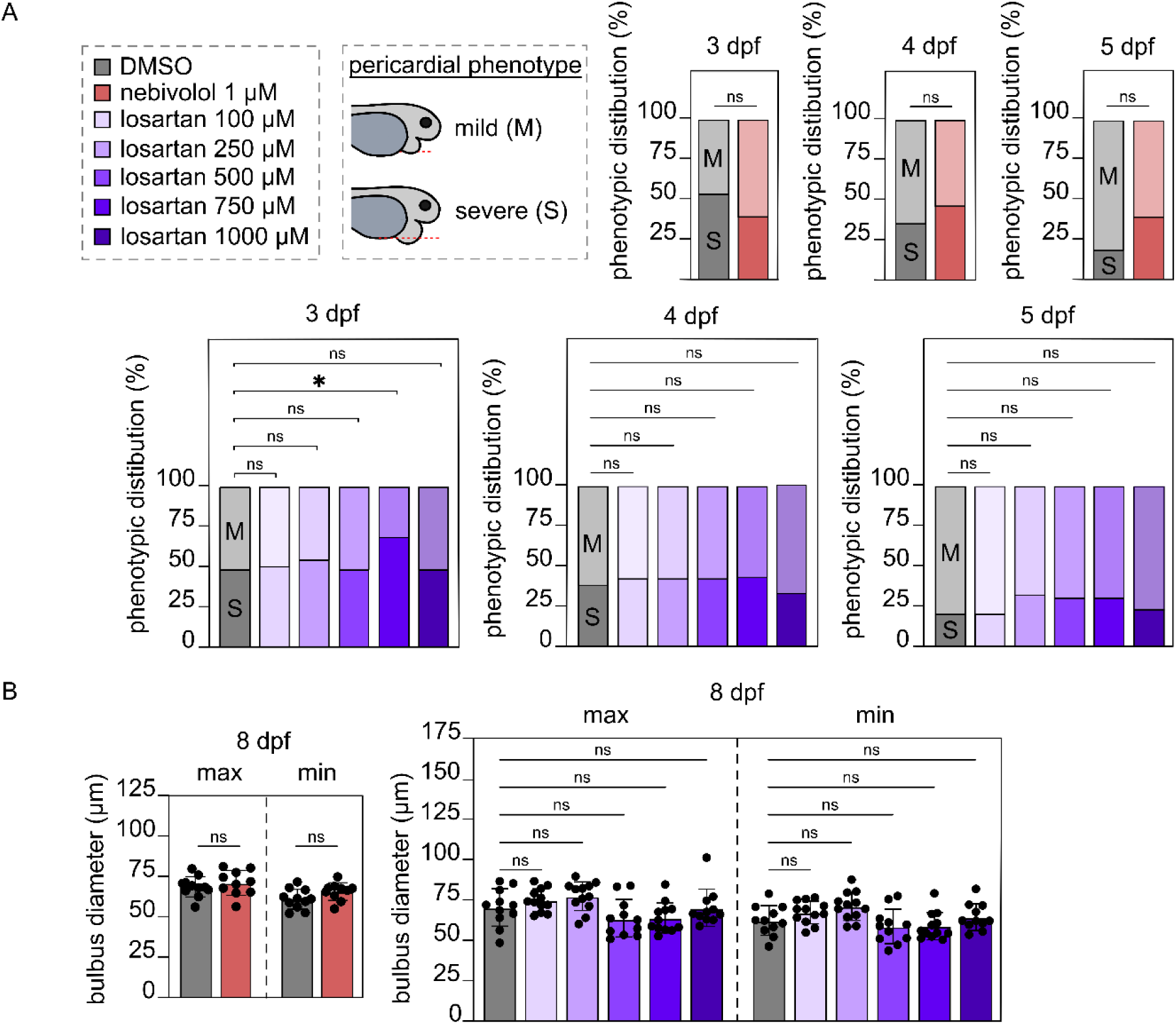
Effects of angiotensin II receptor blocker and β-adrenergic receptor blocker in the fbn2b^-/-^ zebrafish model. **(A)** 1 dpf *fbn2b^-/-^* zebrafish embryos were treated with 100-1000 µM losartan, an angiotensin II receptor blocker (purple-blue shading, n = 12 – 15), and 1 µM nebivolol (red), a β-adrenergic blocker, with solvent as a control (DMSO – grey, n = 70). Distribution of the pericardial phenotype is shown at 3, 4, and 5 dpf. **(B)** At 8 dpf, the diameter of BA was measured, both in maximal (left) and minimal distension (right). Data are expressed as mean ± SEM. ns = non-significant. Statistical test analysis: Kruskal-Wallis test with Dunn’s multiple comparisons test (A), Fisher’s exact test (B). A = atrium, BA = bulbus arteriosus, M = mild pericardial phenotype, S = severe pericardial phenotype.

## DISCUSSION

Due to sustained efforts in translational research, the life expectancy of individuals with MFS has improved dramatically over the past three decades—an achievement that stands out in the field of cardiovascular medicine.^26^ This progress has been driven by advances in diagnostics, longitudinal care, and medical as well as surgical interventions, some of which are underpinned by mechanistic insights gained from mouse models.^27^ These models have been instrumental in identifying key pathways involved in aortic disease progression and in testing therapeutic strategies. Yet, despite these successes, fundamental questions remain unresolved, particularly concerning early developmental processes in MFS, tissue-specific dynamics, functional effects of genetic variants, and targetable disease mechanisms. There is a growing need for complementary model systems that can capture aspects inaccessible in murine models and that allow higher-throughput, *in vivo* functional studies in a vertebrate context. We believe that these issues can be addressed by the new zebrafish model presented in this study, which demonstrated that disruption of *fbn2b* in zebrafish results in a consistent, early-onset Marfan syndrome-like phenotype, including aortic dilatation, valvular defects, and arrhythmia. To the best of our knowledge, this study is the first to investigate the functional orthology between human and zebrafish fibrillins *in vivo*, establishing a foundation for utilizing zebrafish to model MFS and to broaden our understanding of other human fibrillinopathies.

### Zebrafish fbn1 deficiency: a critical look at cardiovascular phenotype reproducibility

The high level of conservation between zebrafish and human fibrillin genes/proteins implies a conserved biological function. However, our findings suggest that the zebrafish *fbn1* isoform may not be the direct functional ortholog of human *FBN1*, as loss of *fbn1* did not result in a cardiovascular phenotype. These results stand in contrast to earlier studies that described mild cardiovascular abnormalities in *fbn1*-deficient zebrafish.^28–30^ Specifically, the study by Chen et al. made a morpholino knockdown of *fbn1* and observed abnormal vessel dilatation in the head, and a lack of venous plexus remodeling.^28^ However, these effects were dose-dependent, with significant abnormalities only seen at relatively high morpholino concentrations. This raises concerns about potential off-target effects and dose-dependent toxicity.^31^ Another study by Yin et al. using heterozygous CRISPR-generated *fbn1^+/-^* mutants with frameshift indels in exon 40 (which is identified as exon 16 in their manuscript, as an incomplete *fbn1* gene sequence was used for reference) showed limited data suggesting phenotypic alterations including body elongation, reduced pigmentation, and altered cardiac blood flow.^29^ More recently, an independent group modeled a likely pathogenic nonsense variant identified in a consanguineous family, by CRISPR/Cas9-based indel generation in exon 60 of *fbn1* (labeled as exon 19 in the study, as, again, an incomplete zebrafish *fbn1* sequence was used). The authors reported a range of phenotypes in their *fbn1^+/-^* mutants, including pericardial edema, tail curvature, and aortic arch bleeding, but no quantitative data was presented showing the frequency of these phenotypes. Other cardiovascular phenotypes described in this study included decreased cardiac function, angiogenesis abnormalities, and increased dorsal aortic diameter, although these phenotypes were not clearly reported.^30^

We were unable to replicate these results in repeated rigorous experiments with any of our four independently generated *fbn1* mutant zebrafish lines. Considering the known influence of genetic background on the cardiovascular severity phenotype in mouse models of MFS,^32^ it is conceivable that differences in zebrafish strains used in these studies (TU versus AB in our study) may partly explain the divergent outcomes of *fbn1* deficiency. However, it is unlikely that strain differences alone account for all observed inconsistencies. We therefore encourage further independent investigations to resolve these discrepancies.

### Lack of compensatory mechanisms underscores fibrillin gene complexity in zebrafish

Another potential reason why complete loss of *fbn1* in our models does not lead to a reproducible phenotype could be that compensatory mechanisms are activated. It has previously been shown that CRISPR-induced frameshift mutations leading to premature stop codons can lead to transcriptional adaptation by increasing the expression of genes with a substantial sequence similarity.^33^ However, while our qPCR analysis indicated a clear downregulation of *fbn1* expression, signifying nonsense-mediated decay, we did not observe compensatory upregulation of other fibrillin genes. Therefore, we hypothesize that the lack of any detectable cardiovascular phenotype in our *fbn1^-/-^* zebrafish rather underscores the complexity of the interspecies functional orthology within the fibrillin gene family.

### Lower homology of zebrafish fbn2a to human fibrillins

We also investigated the cardiovascular impact of *fbn2a* deficiency, both independently and in combination with the loss of *fbn1*, but again did not observe any abnormalities. This result is consistent with the lower amino acid homology of *fbn2a* with human fibrillin genes. Interestingly, fibrillin-2a lacks all RGD motifs, which are known to mediate binding affinity towards various integrins,^34^ while it does contain a novel proline/glutamine-rich domain not found in any other fibrillins. These unique structural differences likely reflect a greater evolutionary distance from the other fibrillins and suggest a unique functional role for zebrafish fibrillin-2a, which may be less relevant to humans.

### In contrast to fbn1 and fbn2a: fbn2b loss leads to developmental defects

Unlike loss of *fbn1* and/or *fbn2a*, disruption of the *fbn2b* gene consistently resulted in an early and reproducible cardiovascular phenotype reminiscent of MFS, persisting into adulthood. This finding suggests that *fbn2b* plays a prominent role during early zebrafish development, functionally resembling human *FBN2*, although the effects during early development have a lasting impact throughout the zebrafish’s lifespan. This notion aligns well with our qPCR data, which demonstrate that expression of *fbn2b* precedes *fbn1* during zebrafish embryogenesis, similar to *FBN2* and *Fbn2* in humans and mice, respectively, suggesting a conserved role for *fbn2b* in early developmental processes.^35^

### CRISPR-generated fbn2b mutant: a phenotypic comparison with existing ENU models

The *fbn2b* mutant zebrafish generated in this study, carrying a premature stop codon in exon 4 (c.458_461del, p.C153*), show a morphological phenotype largely reminiscent of previously reported ENU mutant lines, *puff daddy* (*pfd^gw1^* – p.G161*)^25^ and *scotch tape* (*sco^te382^ – p.C1312F*)^36^. These ENU mutants have either nonsense or missense mutations, resulting in either a loss of protein or an altered protein structure, respectively. Similar to the *pfd^gw1^* and *sco^te382^* mutants, our *fbn2b* homozygous mutants display a characteristic finfold atrophy, which does not appear to impact their overall fitness. In contrast to the *pfd^gw1^* mutant, our CRISPR-induced *fbn2b* mutant line did not develop a notochord phenotype, which was also not present in the *sco^te382^ mutant*. It was later discovered that, besides mutations in *fbn2b*, mutations in the *pku300* gene also contribute to the cardiovascular defects in the *sco^te382^* mutant.^36^ Similarly, it cannot be excluded that confounding genetic factors also influence the observed notochord deformities in *pfd^gw1^*. This might explain why our novel CRISPR/Cas9-generated *fbn2b* mutant model with a frameshift mutation in almost the same location as *pfd^gw1^* does not present with a notochord phenotype. This underscores that, while forward genetic screens are valuable in identifying novel genes within developmental pathways, thorough validation is necessary to exclude the possibility that multiple genes contribute to the observed phenotype.

Consistent with observations in the *sco^te382^* mutant (but unlike *pfd^gw1^*), the severe pericardial edema phenotype was not completely penetrant in our *fbn2b^-/-^* zebrafish. We hypothesize that the progression to irreversible, lethal endocardial detachment is a stochastic process. As described previously for the *sco^te382^* mutant^25^, all *fbn2b^-/-^* zebrafish likely experience compromised endocardial integrity, leading to increased intercellular gap formation. Only a subset reaches a critical threshold in the size of these gaps, leading to the complete detachment of the endocardium from the underlying cardiac jelly.^36^ Interestingly, a subset of our CRISPR-induced *fbn2b^-/-^* mutants initially develops pericardial edema around 2 dpf, which appears to resolve by 5 dpf in approximately 30% of the cases. We hypothesize that in these *fbn2b^-/-^*zebrafish the enlarged endocardial gaps, which are formed dynamically during growth of the cardiac chambers in early development and result in fluid leak into the pericardium, are closed sufficiently during further cardiac developmental stages in these embryos, preventing complete endocardial detachment, restoring the integrity of the endocardial barrier function and ensuring survival.^25^

### BA dilatation in fbn2b mutants reflects aortic pathology seen in MFS

The *fbn2b^-/-^* zebrafish without a pericardial phenotype at 5 dpf survive to adulthood and are fertile. Interestingly, we found that these zebrafish show a dilated BA phenotype that emerges during the larval stage and persists into adulthood. The specific dilation of the BA observed in our *fbn2b^-/-^* model is particularly relevant as the BA is anatomically and functionally related to the aortic root and ascending aorta in humans,^37,38^ which is typically dilated in patients with MFS.^39^ The elastic outflow properties of the BA are evolutionarily conserved, supporting its role as a ‘Windkessel’ organ. Like the aortic root, the BA functions as an elastic reservoir, absorbing the pulsatile output of the ventricle and ensuring continuous blood flow throughout the circulation.^40,41^ Several cases of human *FBN2* defects leading to aortic dilatation have been reported^12–14^, further supporting the indication of a functional overlap between different fibrillins and thus underscoring the potential clinical relevance of this zebrafish model. It is important to note that we do not have evidence that BA dilation in our *fbn2b^-/-^*zebrafish progresses to dissection and fatal aortic rupture at later stages. The only zebrafish line so far known to develop progressive ventral aortic dissection and rupture is a model with defects in the TGF-β effector proteins Smad3 and Smad6.^42^

Dilation of the BA was also observed in previously generated zebrafish mutant lines deficient in various mediators of the TGF-β signaling pathway. Zebrafish lacking both *Tgfbr1* paralogues (*alk5a^-/-^, alk5b^-/-^*) show a severe dilation of the developing BA and ruptures in its endocardial layer starting at 4 dpf, which results in severe pericardial edema, retrograde blood flow, and ultimately mortality by 7 dpf.^43^ Zebrafish double mutant for the latent TGF-β-binding proteins 1 (ltbp1) and 3 (ltbp3) (*ltbp1^-/-^*, *ltbp3^-/-^*) exhibit BA and ventricular dilation during larval stages, pericardial edema, aortic regurgitation, and premature death by 8 dpf.^44^ These phenotypes are similar to the *fbn2b^-/-^*larvae with a severe phenotype, which also die by 8 dpf. These similarities highlight the relevance of fibrillin and TGF-β interactions, and the consequences of their disruption, which are central to the pathogenesis of MFS.^44^

### fbn2b deficiency induces elevated heart rate and arrhythmias

We also report a slightly elevated heart rate in our *fbn2b^-/-^*embryos compared to their WT siblings. Although common MFS mouse models (e.g. *Fbn1^C1041G+^*) are not known to show significant differences in heart rate^45^, a study in human induced pluripotent stem cell-derived cardiomyocytes showed that cells carrying a pathogenic *FBN1* variant had a higher intrinsic beating rate.^46^ Of note, zebrafish cardiac electrophysiology differs significantly from that of mice and can even be considered physiologically more similar to humans.^47,48^ In MFS patients, cardiac arrhythmias, frequently associated with mitral valve prolapse, are well-recognized clinical features^49^, which have been replicated in mouse models.^50^ Our adult *fbn2b^-/-^* zebrafish model recapitulates this aspect, as they exhibit a higher frequency of irregular or skipped heartbeats compared to WT siblings. Given the clinical relevance of cardiac arrhythmias in MFS,^49,51,52^ our *fbn2b^-/-^* zebrafish model can be valuable for further elucidating the underlying physiological mechanisms involved. Interestingly, *fbn2b^-/-^* larvae had a stronger negative chronotropic effect upon administration of the β-adrenergic receptor blocker nebivolol. This observation suggests that *fbn2b^-/-^* zebrafish may have an intrinsically elevated sympathetic tone, which warrants further investigation.

### fbn2b’s role in endothelial mechanosensing

Similar to observations in the *pfd^gw1^* mutant,^25^ we found that the posterior caudal vein in our *fbn2b^-/-^* zebrafish initially develops as a dilated structure with compromised vessel integrity. This phenotype resolves during further development in *fbn2b^-/-^*without persistent endocardial detachment, unless cardiac contraction is pharmacologically inhibited, resulting in loss of blood flow. This suggests that fibrillin mediates blood flow-induced biomechanical signaling crucial for vascular development, aligning with fibrillin’s established role in mechanosensing.^53,54^ In MFS mice (*Fbn1^C1041/G+^*), it has been shown that morphological defects in endothelial cells are most pronounced at aneurysm-prone regions exposed to elevated mechanical stress.^55,56^ Since vascular smooth muscle cells are not recruited to the zebrafish arterial wall until after 3 dpf, the early vascular defects observed here are likely the result of endothelial dysfunction.^57^

### Total fibrillin deficiency in zebrafish

TKO zebrafish, lacking any functional fibrillin gene, showed cardiovascular phenotypes during early development, which are similar to the single *fbn2b* KO. Furthermore, they showed decreased general fitness as indicated by a shorter standard length and slightly decreased survival during larval stages. Interestingly, TKO do not survive to adulthood, suggesting that the additional loss of *fbn2b* unmasked essential functions of *fbn1* and *fbn2a*. This also implies that loss of only one or two zebrafish fibrillins can generally be compensated by the other fibrillin(s) to guarantee (at least partial) survival. However, the essential role of *fbn2b* in cardiovascular homeostasis can not be compensated by the other fibrillins. In mice, *Fbn1* functionally compensates for *Fbn2* loss during embryogenesis in a dosage-sensitive manner. Nearly all *Fbn1^-/-^;Fbn2^-/-^*embryos and approximately half of *Fbn1^+/-^;Fbn2^-/-^* embryos fail to survive gestation, underscoring the critical role of *Fbn1* dosage in development.^58^

### fbn2b deficiency triggers the complement pathway

Bulk RNA sequencing analysis of our *fbn2b* mutant model revealed upregulation of several components of the complement signalling pathway during early developmental stages. The complement system is a key component of innate immunity, mediating host defense through a cascade of proteolytic activations that enhance pathogen clearance and modulate inflammation. It can be activated via three distinct pathways: the classical, lectin, and alternative pathway. Increasing evidence implicates the alternative pathway in the pathogenesis of cardiovascular diseases, including aortic aneurysms. For example, activation of the alternative complement pathway contributes to elastase-induced abdominal aortic aneurysm formation, and studies have shown elevated plasma levels of C3a and C5a in patients with TAAD.^59,60^ Interestingly, genetic ablation of *Cfb* in *Fbn1^C1041/G+^* MFS mice reduced the occurrence of TAAD, further supporting a pathogenic role for this pathway. In a related study, the same group demonstrated the C3a-C3aR axis contributes to TAAD via upregulation of MMP2, a key enzyme involved in ECM remodeling.^60^ Additionally, genetic variants in C1R have been associated with TAA formation in patients with bicuspid aortic valve, suggesting a broader involvement of the complement system in aortic disease.^61^ Collectively, these findings and our current data imply that complement activation may play a role in aneurysm progression in MFS. It is well established that endothelial cells in MFS mice exhibit structural and functional abnormalities that result in endothelial dysfunction,^56^ which is also evidenced in our *fbn2b^-/-^* zebrafish model by the endocardial and caudal vein phenotypes. We propose that these structural defects may serve as triggers for complement activation, leading to the recruitment of inflammatory cells such as macrophages and neutrophils. These cells are known to secrete matrix metalloproteinase-9 (MMP9), and to a lesser extent MMP13, which contributes to extracellular matrix degradation and disease progression in TAAD, particularly by digesting different collagen types, including the main fibrillar and basement membrane collagens. Interestingly, both *mmp9*, *mmp13a* and *mmp13b* were increased in the *fbn2b^-/-^* RNA sequencing dataset, and could represent downstream targets of immune cell activation, which contribute to aortic wall damage. Targeting the complement cascade - particularly components of the alternative pathway - may therefore represent a promising therapeutic strategy for preventing or mitigating TAAD in MFS. However, further studies are required to determine the most effective and specific targets within the complement system.

Collectively, these findings underscore the complex nature of aortic disease in MFS, demonstrating the contribution of various signaling pathways beyond TGF-ß. A deeper understanding of the intricate crosstalk among these pathways is therefore essential and warrants further investigation.

### Cardiac valve abnormalities in adult fbn2b mutants: parallels with patients with MFS

Histological staining for elastin in sections of adult *fbn2b^-/-^* zebrafish revealed cardiac valve abnormalities in both the AV and particularly the BV valves. Propagation-based phase-contrast synchrotron imaging confirmed longer and thicker BV valve leaflets in all *fbn2b^-/-^*, which was supported by the observation of widened valve openings *in vivo* using CFD. Similar cardiac valve defects have been shown in zebrafish lacking the gene coding for elastin isoform a (*elna^sa12235^*).^62^ The phenotypic resemblance between these two zebrafish models suggests a possible shared etiology, potentially reflecting the essential role of fibrillin as a scaffold for elastin fiber formation. On the other hand, it is tempting to speculate that the important role of *fbn2b* in maintaining an intact endocardial barrier through intercellular contact during early development^25^ could be related to the observed valve defects in the adult *fbn2b^-/-^* zebrafish. It has been shown that endocardial cell migration significantly contributes to the initial development of the AV valve in early larval stages.^63^ Additional studies will be needed to confirm whether loss of *fbn2b* affects these processes in the development of both the AV and BV valves, which could explain the malformation of valvular leaflets we observed in adult *fbn2b^-/-^* zebrafish.

Cardiac valve architecture has been closely examined in *Fbn1*^C1041G/+^ and *Fbn1*^C1041G/C1041G^ MFS mouse models as well, both of which show dramatic alterations in mitral valve structure. Notably, *Fbn1*^C1041G/C1041G^ mice display unique malformations where leaflet tips fold back and fuse to more proximal segments,^64,65^ resembling the abnormalities we observed in the *fbn2b^-/-^* zebrafish with an AV valve phenotype.

The valve defects identified in our *fbn2b^-/-^* zebrafish model further support its validity as a model of the cardiovascular manifestations of MFS. Mitral valve prolapse and tricuspid valve prolapse respectively affect approximately 65% and 35% of patients with MFS.^66^ Nevertheless, while valve defects are associated with regurgitation in approximately 40% of cases, this was only observed in a few cases using PWD in our zebrafish model. It is conceivable that the mild valve defects that we observed using histology and 3D modeling may have caused only subtle or intermittent regurgitation that eluded detection by our echocardiography measurements, particularly given the challenges of using this technique in small organisms like zebrafish.^67^ Though a subset of patients with MFS develops intrinsic cardiomyopathy characterized by left ventricular dilation and dysfunction, myocardial dysfunction is at most very mild in the majority of individuals.^68–71^ Echocardiography measurements revealed mild ventricular hypertrophy in our *fbn2b^-/-^*zebrafish, while no statistically significant differences were observed between fibrillin mutants and WT zebrafish in any of the 20 PWD parameters investigated. Synchrotron-based 3D reconstructions of the atrium, ventricle, and BA in *fbn2b^-/-^* zebrafish did not show any significant differences in volume and diameter compared to WT. It is important to note, however, that synchrotron imaging is performed on fixed samples collected postmortem, where cardiac and vascular structures collapse due to loss of intraluminal blood pressure. This discrepancy could potentially be overcome by vascular corrosion casting.^72,73^ Interestingly, *fbn2b^-/-^* zebrafish show significantly higher variability in atrial volumes compared to WT. Those mutants with the largest atria also exhibit pronounced thickening of the AV valves, suggesting a pathophysiological consequence of prolonged volume overload due to the retrograde blood flow across the valve.^74^ However, there were only a limited number of these cases, and echocardiography data is not available for these samples, limiting confirmation.

### Exploring the efficacy of beta-blockers and angiotensin receptor blockers in fbn2b zebrafish

Given that β-adrenergic antagonists are considered standard medical care for MFS patients,^75^ we investigated their effects in our zebrafish model. We tested atenolol, the most prescribed β-blocker in MFS with selective anti-β1-adrenergic receptor activity, and nebivolol, a third-generation, also β1-selective β-blocker but with additional vasodilatory effects mediated by nitric oxide release, which has also been tested in MFS patients.^76,77^ We found that nebivolol significantly decreased heart rate in WT and, to an even larger extent, *fbn2b^-/-^* larvae. Atenolol, however, did not significantly impact heart rate in either genotype. This disparity in the effects of both β-blockers could potentially be explained by differences in pharmacokinetics – it has been shown previously that atenolol has no significant effect on heart rate in zebrafish larvae over a range of doses, while other β-blockers do.^78^ Alternatively, it is conceivable that the zebrafish larvae may be more sensitive to modulation by nitric oxide released after nebivolol administration.

Next, we tested the effects of nebivolol at the dose that was shown to be bioactive based on the heart rate reduction, and losartan, an angiotensin II type I receptor blocker, another commonly prescribed medication for patients with MFS.^79^ We found that neither losartan nor nebivolol had a measurable impact on the *fbn2b^-/-^* pericardial edema phenotype distribution or BA dilation. This result however aligns with clinical observations, showing that β-blockers and angiotensin receptor blockers can slow the rate of increase of aortic dilation in patients with MFS, resulting in a delay in the need for prophylactic aortic surgery, but do not abolish the risk of aortic dissection and rupture, which is unfortunately still encountered in patients under close medical management.^77^ The findings in our zebrafish model imply that these drugs do not directly address the basic pathophysiological mechanisms leading to the cardinal cardiovascular manifestations of MFS. This further underscores the need for novel therapeutic strategies to eliminate the risk of potentially fatal aortic complications in MFS, which can now be addressed using previously unavailable unbiased approaches in our novel zebrafish model.

## STUDY LIMITATIONS

An inevitable limitation of this study concerns the interspecies relationship between the fibrillin genes in human and zebrafish. There is no definitive explanation why loss of zebrafish *fbn2b*, rather than *fbn1*, gives rise to the observed MFS-like phenotypes, while *FBN1* is clearly the primary disease-causing gene in humans. Our data indicate that *fbn2b* is more important during early development, resulting in the observed phenotypes in larvae, which persist until adulthood. The lack of any (progressive) phenotype in zebrafish lacking *fbn1* suggests a compensation by the other fibrillin(s), which is supported by the lack of viability of zebrafish lacking all fibrillin genes. Further studies will, however, be necessary to completely unravel the functional distinctions between the different zebrafish fibrillins. Nevertheless, our data are concordant in showing that loss of *fbn2b* leads to phenotypes that faithfully mimic the cardiovascular manifestations of MFS.

The limitation of our *fbn2b^-/-^* zebrafish model is the observed phenotypic heterogeneity during larval stages. A conclusive explanation of the variable outcome of the endocardial phenotype in *fbn2b^-/-^*larvae, with only a subset developing a lethal phenotype, remains lacking. Our hypothesis that a stochastic process is at the basis of the fate of the potential progression to complete endocardial detachment can only be proven by exclusion of other possible explanations. Finally, in this study, we focused predominantly on cardiovascular manifestations in our zebrafish model, as those are the chief cause of morbidity and mortality in patients with MFS. Additional studies will be necessary to investigate skeletal and ocular manifestations, which are common features in patients with MFS.

## CONCLUSIONS

Our comprehensive study of zebrafish fibrillins demonstrates that disruption of zebrafish *fbn2b* leads to a spectrum of cardiovascular abnormalities that resemble those observed in human MFS, including BA dilation, cardiac valve defects, and heart rhythm irregularities. These findings highlight the complex interspecies orthology of fibrillins. Our data further suggest that the early phenotype observed in our model is potentially related to the activation of the innate immune system. The availability of this novel zebrafish model represents a significant new tool for research into the mechanisms of MFS-related cardiovascular manifestations and the identification of new therapeutic targets.

## CLINICAL PERSPECTIVES

### COMPETENCY IN MEDICAL KNOWLEDGE

Marfan syndrome (MFS) is a life-threatening connective tissue disorder with no effective cure. Understanding of its pathogenesis and available treatment options remains limited, highlighting the urgent need for more versatile animal models. We show that zebrafish with *fbn2b* deficiency exhibit cardiovascular abnormalities that closely resemble those observed in patients with MFS. Our data suggest that fibrillin defects disturb essential cardiovascular development processes, which is not addressed by currently recommended medical treatments for patients with MFS.

### TRANSLATIONAL OUTLOOK

The availability of a zebrafish model that faithfully recapitulates multiple cardiovascular manifestations of MFS enables novel strategies to pursue better diagnosis and treatment options for patients. The cardiovascular phenotype of *fbn2b*^⁻/⁻^ zebrafish larvae can serve as a valuable readout for unbiased *in vivo* high-throughput drug screening to uncover new therapeutic targets. The model may also be of value to test the pathogenicity of rare *FBN1* variants identified with diagnostic testing.

### HIGHLIGHTS

- We disrupted zebrafish fibrillins to create a novel animal model of MFS.
- We applied detailed cardiovascular phenotyping across both early developmental and adult stages of fibrillin-impaired zebrafish.
- *fbn2b^-/-^* zebrafish exhibit aortic dilation and valvular abnormalities resembling the cardiovascular manifestations observed in patients with MFS.
- This *fbn2b^-/-^* zebrafish model represents a relevant tool for advancing our understanding of MFS pathogenesis and for identifying urgently needed therapeutic strategies.

## ACKNOWLEDGEMENTS

We wish to thank the Zebrafish Facility Ghent Core, particularly Karen Vermeulen for the diligent care of our zebrafish. We also thank our NGS core and Ghent Light Microscopy Core. We acknowledge Diamond Light Source (Harwell Science and Innovation Campus, UK) for time on Beamline I13-2 under Proposal MG32919-1, and Swiss Light Source (Paul Scherrer Institut, Switzerland) for time on beamline TOMCAT X02DA under Proposal 20221979. We would like to thank Violette Deleeuw, Michiel Vanhooydonck, Yousof Mohammad Asaad Abdel-Raouf, Simon D’hulst and Isaac Rodriguez-Rovira for their assistance in acquiring synchrotron data. We acknowledge the use of AI tools (ChatGPT, Copilot and Gemini) for text refinement.

## ABBREVIATIONS LIST

AV: atrioventricular (valve)
BA: bulbus arteriosus
BV: bulboventricular (valve)
CFD: color flow Doppler
dpf/hpf/mpf: days/hours/months post fertilization
TKO/KO: triple/knock-out
MFS: Marfan syndrome
PWD: pulsed-wave Doppler
TAAD: thoracic aortic aneurysm and dissection
TGF-β: transforming growth factor-β
WT: wild type

