## Supplemental Appendix for "Systematic disruption of zebrafish fibrillin genes identifies a translational zebrafish model for Marfan syndrome"

### **Supplemental Methods**

**Zebrafish husbandry**

Zebrafish were maintained in a semi-closed recirculating housing system (ZebTec, Tecniplast, Buguggiatte, Italy) at a constant temperature (27–28 °C), pH (~7.5), conductivity (~550 mS), and a light/dark cycle (14 h/10 h). Fish were fed twice daily with dry food (Gemma Micro, Skretting, Stavanger, Norway) and once with Micro Artemia (Ocean Nutrition, Essen, Belgium). Breeding and the collecting of the embryos were performed according to previously described protocols.^1^ All experiments were performed on offspring from stable mutant lines which were backcrossed for at least four generations to WT zebrafish to avoid confounding off-target effects. All experiments were approved by the Animal Ethics Committee of the Ghent University Faculty of Medicine and Health Sciences (ECD 17-75K, ECD 17-78 and ECD 19-16K) and conform to the guidelines from Directive 2010/63/EU of the European Parliament on the protection of animals used for scientific purposes. All efforts were made to minimize pain, distress, and discomfort.

**Generation of fibrillin mutant zebrafish lines**

Single Guide RNA (sgRNA) sequences used for the generation of the various mutant fibrillin lines are summarized in Supplemental Table 1. gBlocks containing the entire sgRNA sequence preceded by a T7 promotor were obtained from Integrated DNA Technologies (Leuven, Belgium), and used as template for in vitro transcription using the MEGAshortscript™ Kit (Thermo Fisher Scientific, Waltham MA, USA) according to the manufacturer’s instructions. Purified sgRNA transcripts were then injected together with Cas9 protein (Toolgen, Seoul, Republic of Korea) as a ribonucleoprotein complex into fertilised eggs of zebrafish carrying the *Tg(kdrl:EGFP)* reporter at the single-cell stage using a Femtojet 4i microinjector (Eppendorf, USA). Embryos were maintained in an incubator (28 °C) in E3 medium until 5 dpf prior to transfer into the zebrafish facility. Genetically stable mutant zebrafish lines were established by backcrossing with wild type (WT, AB background) zebrafish.

The genotype of zebrafish was determined by finclipping the tail at 3 months post fertilization (mpf). Fish were shortly anesthetized using 1x tricaine methanesulfonate solution. Briefly, DNA was extracted by heating the caudal fin biopsy in 100 ml 50 mM sodium hydroxide (NaOH) at 95 °C for 20 minutes, followed by the addition of 10 ml 1 M Tris-HCl (pH 8) to neutralize the solution before this was added to the PCR reaction mix. The primer pairs used for PCR amplification and Sanger sequencing are summarized in Supplemental Table 2. Thermocycling conditions were 95 °C for 3 minutes, followed by 35 cycles of 95 °C for 15 seconds, 58 °C for 10 seconds, and 72 °C for 15 seconds, and a final extension of 72 °C for 10 minutes. Finally, Sanger sequencing was performed on amplified fragments, using the same primer set as for the PCR amplification.

**RNA extraction for RNAseq and RT-qPCR**

Total RNA of 10 pooled WT, heterozygous and/or homozygous mutant *fbn2b* zebrafish, collected at specific timepoints as mentioned in the corresponding figure legends, was extracted using Trizol^ã^ (Life Technologies Europe). Subsequently, RNA was purified using the RNeasy Mini Kit (Qiagen, Hilden, Germany) in combination with on-column DNAse I treatment (Qiagen, Hilden, Germany) according to the manufacturer’s guidelines. RNA concentration and purity was quantified using the Little Lunatic (Unchained Labs, Pleasanton, CA, USA) and RNA quality was determined using Tapestation (Agilent, USA) according to the manufacturer’s guidelines. Only samples with RNA integrity number (RIN) values higher than 9 were considered. Next-generation sequencing libraries were prepared using the TruSeq Stranded Total RNA kit (Illumina, USA) following the manufacturer’s recommended procedures. Sequencing was then performed on an a NovaSeq6000 instrument (Illumina, USA). Following quality control,^2^ raw reads were aligned to the reference genome GRCz11 using STAR.^3^ Data normalization and filtering were conducted using standard workflows in R.^4–6^ Differential expression analysis was performed with edgeR, applying corrections for multiple testing.^7,8^ Differentially expressed genes (DEGs) were defined using a false discovery rate (FDR)-adjusted p-value threshold of 0.05 and a minimum fold change of 1. GO enrichment analysis was subsequently performed to identify affected biological pathways.^9^

cDNA was synthesized starting from 750 ng RNA using the iScript cDNA Synthesis Kit (Bio-Rad Laboratories, Hercules, CA, USA), and primers (IDT; Integrated DNA Technologies) were designed using IDT’s PrimerQuest tool. For each cDNA sample, RT-qPCR reactions were prepared in triplicate with the addition of SsoAdvancedä Universal SYBRâ Green Supermix (Bio-Rad Laboratories, Hercules, CA, USA) prior to analysis on a Roche LightCycler 480 System (Roche Diagnostics, Mannheim, Germany). The reference genes *elfa* and *bactin2* were used for normalization, which primer pairs are shown in Supplemental Table 3.^10^ Cq values were analysed using Excel.

***In vivo* imaging of transgenic zebrafish larvae**

Live WT and mutant zebrafish larvae of various ages were anesthetized and embedded in a glass-bottom WillCo-dish (Willco Wells, Amsterdam, Netherlands) using 1% SeaPlaqueä low melting temperature agarose (Lonza, Basel, Switzerland) dissolved in E3 embryo medium with 0.5x tricaine. The embryos were either positioned with their heart directed towards the bottom of the dish to ensure optimal cardiac and ventral aorta visualisation or sideways positioned for optimal visualisation of the caudal vasculature. Images were captured using an inverted widefield microscope (Zeiss Axio Observer Z1), equipped with a Zeiss Axiocam 503 colour top mounted camera and operated by Zeiss ZenPro software.

To obtain a single in-focus brightfield image of the whole zebrafish larvae, z-stacks of 4 adjacent tiles were first acquired and merged along the optical axis using the Extended Depth of Focus (EDF) plugin in ImageJ software v.1.54g (National Institutes of Health, USA).^11^ The built-in stitching plugin of ImageJ was then used to merge the 4 individual focused tiles along the x-axis. Fluorescent images were acquired with the same inverted widefield microscope.

Brightfield imaging was used to evaluate cardiovascular function of 3 dpf WT and *fbn2b^-/-^* larvae, as previously described.^12^ Briefly, short recordings (~ 4-5 cardiac cycles) with 60 frames/sec were recorded of larvae in a lateral position. The ventricular area of end-diastole and end-systole from three consecutive cardiac cycles was then used to quantitate different parameters: heart rate (bpm), stroke volume (nl), cardiac output (nl/min), ejection fraction (%), systolic volume (nl) and diastolic volume (nl). Measurements were made with ImageJ analysis software.

For deep tissue imaging, the Nikon A1R HD multiphoton microscope equipped with a Mai Tai DeepSee IR laser (740 – 1040 nm) and a high sensitivity non-descanned detector was used. WT and mutant larvae at different developmental stages were immobilized in low melting temperature agarose, as previously described. To minimize image distortion caused by the beating of the heart, zebrafish were given an overdose of tricaine just prior to acquisition. A maximum intensity projection fluorescence image was created through reconstruction of approximately 170 optical sections of a 0.85 mm z-step, using ImageJ.

**Pharmacological manipulation with myosin inhibitor**

The heartbeat of WT and mutant *fbn2b* larvae was stopped by adding 10 mM myosin inhibitor 2,3-butanedione 2-monoxime to E3 embryo medium. In parallel, control WT and mutant *fbn2b* larvae were exposed to vehicle (0.01% dimethyl sulfoxide (DMSO)). The compounds were added to the petri dish prior to the start of systemic blood circulation (± 22 hours post fertilization (hpf)), followed by manual dechorionation of the embryo. Embryos were embedded sideways in 1% low melting temperature agarose, as mentioned above, with the addition of either 20 ml myosin inhibitor or DMSO. Z-stack imaging of the caudal vein of 24 and 48 hpf larvae was acquired using the same inverted widefield microscope (Zeiss Axio Observer Z1), equipped with a Zeiss Axiocam 503 colour top mounted camera and operated by Zeiss ZenPro software. ImageJ was used to perform caudal vein measurements.

**Transthoracic echocardiography of adult zebrafish**

Two-dimensional transthoracic echocardiography was performed using a Vevo 2100 ultrasound machine (Fujifilm VisualSonics, Toronto, Canada) and MS 700 linear array probe (50 MHz) (Fujifilm, VisualSonics, Toronto, Canada) according to the detailed protocol by Van Impe, et al.^13^ Briefly, zebrafish were anesthetized in a 1x tricaine solution at 28 °C until a lack of response to external stimuli was observed. Next, zebrafish were stabilised in a customized 3D printed holder between two spongy clamps, submerged with 0.5x tricaine solution at 28 °C to maintain anaesthesia during the experiment, while minimizing the impact of anaesthesia on the cardiovascular parameters. All measurements were obtained within 3 – 4 minutes after the start of anaesthesia induction.

Pulsed-wave Doppler (PWD) and Colour Flow Doppler (CFD) ultrasound recordings were used for defining ventricular and bulbus arteriosus (BA) measurements *in vivo*, as well as quantifying dynamics of blood flow through the atrioventricular (AV) valve (ventricular inflow) and bulboventricular (BV) valve (ventricular outflow). PWD recordings were processed in a completely automated and unbiased manner in MATLAB v23.2.0 (R2023b) (MathWorks, Natick, MA), using previously published and validated algorithms.^13^ Overview of PWD ventricular inflow and outflow parameters is indicated in Supplemental Table 4.

The Matlab script also provided a visual overview of the heartbeat peaks during the 20-second-long recordings. To investigate beat-to-beat variability, therefore checking for heartbeat irregularity, we manually scored the heartbeat recordings. Score 1 to 4 was given to each sample, ranging from score 1 for recordings with no heartbeat variability, to score 4 for samples with highly irregular heartbeats.

Hemodynamic measurements of the ventricle and BA, as well as the analysis of the CFD ultrasound recordings, were manually performed using the Vevo LAB^TM^ analysis software package v5.8.2 (Fujifilm VisualSonics, Toronto, Canada). Tracings of the walls of the ventricle and BA were done in a 2D longitudinal axis view. Ventricular diastolic and systolic parameters are also indicated in Supplemental Table 4. To obtain the maximal dimensions of BA, its tracings were done immediately following BV valve contraction, revealing its maximal area, volume, and diameter.

CFD recordings in an abdominocranial axis view were used for directly visualising blood flow through the valves and possible regurgitation. Ventricular inflow and outflow areas were measured while in maximal intensity and averaged among three cardiac cycles. Regurgitation fractions of each were calculated as the ratios of inflow or outflow areas and their simultaneous retrograde flow.

WT and mutant zebrafish used for the echocardiography experiments ranged from 6 to 16 mpf but were age-matched for each experiment. To account for age- and sex-related differences in body sizes, ventricular and BA volumes were normalized to body surface area (recommended by Wang et al.), which was calculated by the formula body surface area=8.46×weight^0.66^.^14^ The same was performed for PWD parameters atrial wave, atrial regurgitation, aortic wave, and aortic regurgitation, as recommended by Van Impe at al. As it was proven to have significant male-female differences, only zebrafish males were used for calculating the early peak and velocity time interval of the early peak. All of the analyses were done by operators blinded to the genotype.

**Elastic staining of adult zebrafish**

Adult zebrafish were euthanized by lethal dose of tricaine (1 g/L) and fixed in 4% paraformaldehyde in phosphate-buffered saline solution for at least 24 h, then decalcified with citric acid (45% formic acid, 5% sodium citrate) for 4-6 h. The samples were stored in 70% ethanol until paraffin embedding and microtome cutting into 5 μm thick sections using a Leica TP1020 tissue processor. A combination of Weigert’s Hematoxylin and Resorcin-Fuchsin staining was used for distinguishing nuclei and elastic fibers, respectively, with Van Gieson’s counterstain to examine collagen and miscellaneous tissue elements.

**Synchrotron imaging and 3D reconstruction of zebrafish hearts**

Propagation-based phase-contrast synchrotron X-ray imaging was performed at the TOMCAT (X02DA) beamline of the Swiss Light Source (Paul Scherrer Institute, Villigen, Switzerland) and at the I13-2 beamline of Diamond Light Source (Harwell Science and Innovation Campus, Oxfordshire, UK). *fbn2b^-/-^* zebrafish aged 16 and 18 mpf, along with age-matched WT controls, were fixed, decalcified, and stored in 70% ethanol. Directly before scanning, samples were immobilized vertically in 1.5 ml Eppendorf tubes using 1% low- melting temperature agarose. Custom-made sample holders were used to secure the Eppendorf tubes to the rotation stage between the X-ray source and CCD camera.

Image stacks with a resolution of 2560 × 2560 pixels and 2160 slices were captured by a photon detector mounted on a visual light microscope. Total magnification was 4×, with a field of view of 4.16 mm × 4.16 mm (in-plane) × 3.51 mm (axial), and an isotropic voxel size of 1.625 µm^3^. Tomographic reconstruction, including phase retrieval, was performed on-site using Paganin’s algorithm.^15^ Detailed scan parameters of the individual beamlines optimized for zebrafish samples^16^ are provided in the Supplemental Table 5.

Image processing was performed using the medical software package Mimics v25.0 (Materialise, Leuven, Belgium). The zebrafish hearts, including the atrium, ventricle, cardiac valves, and BA with aorta until the first branching site, were semi-automatically segmented. 3D models of the zebrafish hearts were generated from these masks, enabling analysis of volumes or 2D dimensions, such as valve length and thickness or BA diameter, and their comparisons between the genotypes (*fbn2b^-/-^* vs WT).

**Response to β-adrenergic receptor agonist/antagonists**

WT and *fbn2b^-/-^* zebrafish embryos aged 1 or 2 dpf were treated with β-adrenergic receptor antagonists (nebivolol 10 µM, atenolol 10 and 100 µM), and a β-adrenergic receptor agonist (epinephrine 200 µM and 400 µM) by adding the DMSO-dissolved compounds to the E3 medium (final DMSO concentration = 1%). After 24 or 48 hours of incubation, individual zebrafish were placed on a glass slide and imaged under an inverted widefield microscope, focusing on the heart region. Ten-second recordings were made at 60 frames per second.

Heart rate measurement and analysis were done according to the protocol of Gaur *et al 35* and using the formula: heart rate (bpm) = 60 / (average number of frames between heartbeats × time per frame). The change in heart rate after nebivolol treatment was measured as the difference between the mean heart rate in untreated controls and the heart rate in the treated zebrafish, for each sample individually.

**Drug screening in fbn2b^-/-^ zebrafish model using angiotensin II receptor antagonist and β-adrenergic antagonist as commonly used medications in MFS**

*Tg(kdrl:GFP) fbn2b^-/-^* zebrafish embryos aged 1 dpf were treated with losartan (100 µM to 1000 µM), an angiotensin receptor antagonist and nebivolol (1 µM), a β-adrenergic receptor antagonist. Solutions were refreshed daily. The phenotypic distribution of compound-treated *fbn2b^-/-^* zebrafish was evaluated at 3, 4, and 5 dpf, as a percentage of fish with the severe pericardial phenotype in a clutch (n = 70), in comparison to the solvent-treated mutants. Classification of severe and mild phenotypes was done by observing the pericardial edema of a mutant zebrafish – whether its size exceeded the size of the yolk.

At 8 dpf, mutant zebrafish larvae were embedded in 1% low-melting temperature agarose and positioned on their ventral side, as previously described. Fluorescent images of BA were taken for 20 frames (836 ms). Using Zeiss Zen v3.9 software, the diameter of BA was measured, both in maximal and minimal distension. Diameters of compound-treated zebrafish were compared to the solvent-treated mutants.

**Statistical analysis**

GraphPad Prism v10.1.2 software package (GraphPad Software, Boston, USA) was used for all statistical analyses. P-values of the difference in phenotypic distribution were analysed by Fisher’s exact test. All the other P-values were determined using an unpaired two-tailed Student’s t-test or one-way ANOVA with Dunnett’s multiple comparisons test for samples that were normally distributed. Otherwise, the Mann-Whitney, Kruskal-Wallis or Wilcoxon’s signed-rank tests were used. A significance threshold of P ≤ 0.05 was applied, with P ≤ 0.002 when Bonferroni's adjustment was required. All data are presented at the means ±

**Supplemental Table 1.** Sequence of sgRNA used for each genotype.

| **Gene** | **Name** | **Exon** | **Size INDEL** | **sgRNA sequence** | **Predicted**  **nucleotide change** | **Predicted protein change** |
| --- | --- | --- | --- | --- | --- | --- |
| *fbn1* | Cmg79 | 34 | 4bp deletion | GGCTGGACGAGTGCTCGAAC | c.4223_4226del | p.S1408fs |
|  | Cmg57 | 38 | 4bp deletion | GGCCTGGACGTCCAGTACCG | c.4737_4740del | p.Q1579fs |
|  | Cmg64 | 38 | 4bp insertion | GGAACAGCAGCAGGACGCTT | c.4801_4802insTGAT | p.A1601fs |
|  | Cmg80 | 2 | 16bp deletion | GGGTGGAAAACCCTGACCGG | c.259_274del | p.L87fs |
| *fbn2a* | Cmg86 | 5 | 2bp deletion | TGCCACAGTGGGTCAAGCGT | c.581_582del | p.A194V |
|  | Cmg87 | 5 | 22bp deletion | TGCCACAGTGGGTCAAGCGT | c.558_599del | p.L186= |
|  | Cmg95 | 5 | 12bp deletion | TGCCACAGTGGGTCAAGCGT | c.579_590del | p.Q194H |
| *fbn2b* | Cmg96 | 4 | 4bp deletion | TGCGAGAGCGGATGTCAGAA | c.458_461del | p.C153* |
|  | Cmg97 | 4 | 15bp deletion | TGCGAGAGCGGATGTCAGAA | c.462_476del | p.Q154R |

**Supplemental Table 2.** Primer pairs used for genotype verification.

| **Gene** | **Name** | **Exon** | **Size INDEL** | **Forward primer** | **Reverse primer** |
| --- | --- | --- | --- | --- | --- |
| *fbn1* | Cmg79 | 34 | 4bp deletion | ATTAACGAGTGTGAGATCGGG | GGCTTCTACTGCACTGGTAC |
|  | Cmg57 | 38 | 4bp deletion | AGCAATCATAAGCACGAGATCC | CCTGCTGCTGTTCTAGGG |
|  | Cmg64 | 38 | 4bp insertion | AGAATTGTATATGCAGGTCCAG | CCTGGAGGAGAAGGATTCAGA |
|  | Cmg80 | 2 | 16bp deletion | TTTATGAGTGTTTGACCCCTCC | GCTGGAGGAAACAGGATAAGT |
| *fbn2a* | Cmg86 | 5 | 2 bp deletion | ATGTAAATCGCAGACTACCGGA | CCAAGGTCAGCACTGATTTTAT |
|  | Cmg87 | 5 | 22bp deletion | ATGTAAATCGCAGACTACCGGA | CCAAGGTCAGCACTGATTTTAT |
|  | Cmg95 | 5 | 12bp deletion | ATGTAAATCGCAGACTACCGGA | CCAAGGTCAGCACTGATTTTAT |
| *fbn2b* | Cmg96 | 4 | 4bp deletion | TTCTGGGTCACACTGACACG | ACATTGCTTGCTTGGGATATTTGT |
|  | Cmg97 | 4 | 15bp deletion | TTCTGGGTCACACTGACACG | ACATTGCTTGCTTGGGATATTTGT |

**Supplemental Table 3.** Summary of primers used for RT-qPCR.

| **Reference target** | **Forward primer** | **Reverse primer** |
| --- | --- | --- |
| *fbn1* | GTTCAGGCTCCTAAACCCTGT | GACAGTTGTGCTGCTTGGTG |
| *fbn2a* | GAGCAGTGCCCACCTATCAG | CACAGATCGGGGACAACCTT |
| *fbn2b* | TGGTGGGAACATACCAGTGC | AGCTTCCCTCCGAGTTTGTG |
| *elfa* | *GGAGACTGGTGTCCTCAA* | *GGTGCATCTCAACAGACTT* |
| *bactin2* | *TGAGCTGAAACTTTACAGACACAT* | *AGACTTTGGTGTCTCCAGAATG* |

**Supplemental Table 4.** Overview of the cardiovascular parameters obtained by two-dimensional transthoracic echocardiography of

adult fibrillin-impaired zebrafish.

| **Abbreviation** | **Unit** | **Description** | **Method of parameter measurement** |
| --- | --- | --- | --- |
| **A peak** | **mm/s** | peak velocity of the blood flow through the atrioventricular valve during diastole (inflow) | automated PWD |
| **A VTI** | **mm** | velocity time integral of the blood flow through the atrioventricular valve during diastole (inflow) | automated PWD |
| **A regurg** | **mm/s** | peak value of the atrioventricular valve regurgitation | automated PWD |
| **E peak** | **mm/s** | peak velocity of the passive blood flow through the atrioventricular valve during early diastole (inflow) | automated PWD |
| **E VTI** | **mm** | velocity time integral of the passive blood flow through the atrioventricular valve during early diastole (inflow) | automated PWD |
| **AET** | **ms** | aortic ejection time, i.e. the duration of bulboventricular valve being opened | automated PWD |
| **NFT** | **ms** | no flow time, i.e. the duration of atrioventricular valve being closed | automated PWD |
| **HR** | **beat/min** | heart rate | automated PWD |
| **HR st dev** | **-** | standard deviation of heart rate values | automated PWD |
| **Ao peak** | **mm/s** | peak velocity of the blood flow through the bulboventricular valve during systole (outflow) | automated PWD |
| **Ao VTI** | **mm** | velocity time integral of the blood flow through the bulboventricular valve during systole (outflow) | automated PWD |
| **Ao regurg** | **mm/s** | peak value of the aortic valve regurgitation | automated PWD |
| **area;d** | **mm^2^** | ventricular area during diastole | ventricular tracing |
| **area;s** | **mm^2^** | ventricular area during systole | ventricular tracing |
| **CO** | **ml/min** | cardiac output | ventricular tracing |
| **EF** | **%** | ventricular blood ejection fraction | ventricular tracing |
| **FS** | **%** | ventricular fractional shortening | ventricular tracing |
| **SV** | **µl** | ventricular stroke volume | ventricular tracing |

| **volume;d** | **µl** | ventricular volume during diastole | ventricular tracing |
| --- | --- | --- | --- |
| **volume,s** | **µl** | ventricular volume during systole | ventricular tracing |
| **inflow area** | **mm^2^** | amount of blood (in 2D) flowing to the ventricle through the atrioventricular valve during diastole (inflow) | CFD |
| **outflow area** | **mm^2^** | amount of blood (in 2D) flowing to the bulbus arteriosus through the bulboventricular valve during systole (outflow) | CFD |
| **inflow regurg fraction** | **%** | relative amount of blood (in 2D) that returns to the atrium through the atrioventricular valve during diastole; inflow area/retrograde flow area | CFD |
| **outflow regurg fraction** | **%** | relative amount of blood (in 2D) that returns to the ventricle through the bulboventricular valve during systole; outflow area/retrograde flow area | CFD |

**Supplemental Table 5.** Overview of scan parameters across two synchrotron facilities.

| **Synchrotron facility** | Swiss Light Source, Paul Scherrer Institute, Switzerland | Diamond Light Source, Harwell Science and Innovation Campus, UK |
| --- | --- | --- |
| **Beamline** | TOMCAT (X02DA) | I13-2 |
| **Beam energy** | 21.8 keV monochromatic beam | 27 keV pink beam |
| **Object-detector distance** | 250 mm | 495 mm (-415 mm) and 295 mm (-615 mm) |
| **Number of projections** | 1501 | 1501 |
| **Scintillator** | LUAg:CE 20 µm | LuAg 500 µm |
| **Objective** | UPLAPO 4× objective | PlanApoN 2× objective |
| **Camera** | PCO.Edge 5.5 | PCO.Edge 5.5 |

**Supplemental Table 6.** P-values of cardiovascular parameters obtained by two-dimensional transthoracic echocardiography of adult *fbn1* and *fbn1*, *fbn2a* double mutant zebrafish, compared to age-matched WT or fbn1 mutants, respectively.

|  | ***fbn1*** | | | | ***fbn1, fbn2a*** | | | |
| --- | --- | --- | --- | --- | --- | --- | --- | --- |
|  | ***fbn1 cmg57*** | ***fbn1 cmg64*** | ***fbn1 cmg79*** | ***fbn1 cmg80*** | ***fbn1 cmg80, fbn2a cmg86*** | ***fbn1 cmg80, fbn2a cmg87*** | ***fbn1 cmg80, fbn2a cmg95*** | ***fbn1 cmg79, fbn2a cmg95*** |
| **A peak** | 0.31 | 0.88 | 0.58 | 0.020 | 0.63 | 0.026 | 0.22 | 0.57 |
| **A VTI** | 0.021 | 0.43 | 0.85 | 0.57 | 0.25 | 0.045 | 0.40 | 0.82 |
| **A regurg** | 0.23 | 0.46 | 0.56 | 0.082 | 0.96 | 0.72 | 0.86 | 0.057 |
| **E peak** | 0.85 | 0.20 | 0.67 | 0.071 | 0.39 | 0.083 | 0.98 | 0.30 |
| **E VTI** | 0.31 | 0.13 | 0.91 | 0.79 | 0.25 | 0.11 | 0.76 | 0.41 |
| **AET** | 0.96 | 0.93 | 0.89 | 0.30 | 0.69 | 0.40 | 0.42 | 0.087 |
| **NFT** | 0.13 | 0.85 | 0.21 | 0.33 | 0.70 | 0.31 | 0.27 | 0.47 |
| **HR** | 0.34 | 0.75 | 0.22 | 0.61 | 0.93 | 0.15 | 0.74 | 0.70 |
| **HR st dev** | 0.88 | 0.23 | 0.59 | 0.75 | 0.57 | 0.67 | 0.53 | 0.61 |
| **Ao peak** | 0.67 | 0.24 | 0.29 | 0.51 | 0.36 | 0.69 | 0.24 | 0.73 |
| **Ao VTI** | 0.52 | 0.055 | 0.46 | 0.96 | 0.74 | 0.46 | 0.95 | 0.51 |
| **Ao regurg** | 0.25 | 0.022 | 0.77 | 0.96 | 0.44 | 0.17 | 0.92 | 0.18 |
| **area;d** | 0.005 | 0.38 | 0.78 | 0.48 | 0.11 | 0.98 | 0.53 | 0.31 |
| **area;s** | 0.012 | 0.24 | 0.47 | 0.37 | 0.49 | 0.97 | 0.61 | 0.87 |
| **CO** | 0.033 | 0.41 | 0.95 | 0.24 | 0.035 | 0.94 | 0.50 | 0.030 |
| **EF** | 0.69 | 0.95 | 0.77 | 0.044 | 0.49 | 0.36 | 0.27 | 0.51 |
| **FS** | 0.41 | 0.16 | 0.63 | 0.63 | 0.72 | 0.92 | 0.081 | 0.35 |
| **SV** | 0.032 | 0.37 | 0.73 | 0.50 | 0.057 | 0.66 | 0.53 | 0.030 |
| **volume;d** | 0.001 (↑) ^a^ | 0.79 | 0.65 | 0.42 | 0.034 | 0.63 | 0.70 | 0.093 |
| **volume,s** | 0.007 | 0.61 | 0.49 | 0.20 | 0.20 | 0.75 | 0.94 | 0.62 |
| **inflow area** | 0.043 | 0.24 | 0.64 | 0.86 | 0.53 | 0.36 | 0.56 | 0.007 |
| **outflow area** | 0.17 | 0.024 | 0.86 | 0.066 | 0.29 | 0.53 | 0.69 | 0.075 |
| **inflow regurg fraction** | 0.50 | 0.40 | 0.78 | 0.62 | 0.94 | 0.46 | 0.000 | 0.85 |
| **outflow regurg fraction** | 0.50 | 0.81 | 0.65 | 0.94 | 0.76 | 0.48 | 0.14 | 0.28 |

No significant differences were observed in any cardiovascular parameters. Statistical analysis: Shapiro-Wilk normality testing, followed by an unpaired t-test was used for normally distributed data, while non-normally distributed data was analyzed using the Mann-Whitney or Wilcoxon signed-rank tests. Bonferroni-adjusted values are reported (significance threshold: P ≤ 0.002), and the direction of change is indicated by an arrow (↑higher or ↓lower). The abbreviations list and explanations of the cardiovascular parameters are included in Supplemental Table 1.

^a^ Higher ventricle volume of the *fbn1 cmg57* mutants has been attributed to be clutch-specific and should not be interpreted as a significant finding, especially when comparing it to the other *fbn1* mutant lines.

**Supplemental Table 7.** P-values of cardiovascular parameters obtained by two-dimensional transthoracic echocardiography of adult *fbn2b* impaired zebrafish, compared to respective age-matched WT.

|  | ***fbn2b*** | |
| --- | --- | --- |
|  | ***fbn2b cmg96*** | ***fbn2b cmg97*** |
| **A peak** | 0.16 | 0.093 |
| **A VTI** | 0.053 | 0.22 |
| **A regurg** | 0.43 | 0.17 |
| **E peak** | 0.007 | 0.48 |
| **E VTI** | 1.00 | 0.15 |
| **AET** | 0.45 | 0.30 |
| **NFT** | 0.93 | 0.036 |
| **HR** | 0.71 | 0.15 |
| **HR st dev** | 0.082 | 0.87 |
| **Ao peak** | 0.89 | 0.34 |
| **Ao VTI** | 0.51 | 0.88 |
| **Ao regurg** | 0.27 | 0.40 |
| **area;d** | 0.054 | 0.68 |
| **area;s** | 0.028 | 0.82 |
| **CO** | 0.98 | 0.44 |
| **EF** | 0.42 | 0.55 |
| **FS** | 0.83 | 0.41 |
| **SV** | 0.70 | 0.44 |
| **volume;d** | 0.076 | 0.54 |
| **volume,s** | 0.024 | 0.70 |
| **inflow area** | 0.0001 (↑) | 0.53 |
| **outflow area** | 0.001 (↑) | 0.001 (↑) |
| **inflow regurg fraction** | 0.45 | 0.063 |
| **outflow regurg fraction** | 0.23 | 0.25 |

The *fbn2b cmg96* mutant line exhibits increased inflow and outflow areas, with higher outflow also observed for the *cmg97* line. These observations are in line with the increase in length and thickness of the valve leaflets revealed by synchrotron imaging. No significant differences were observed in any other cardiovascular parameters compared to WT. Statistical analysis: Shapiro-Wilk normality testing, followed by an unpaired t-test was used for normally distributed data, while non-normally distributed data was analyzed using the Mann-Whitney or Wilcoxon signed-rank tests. Bonferroni-adjusted values are reported (significance threshold: P ≤ 0.002), and the direction of change is indicated by an arrow (↑higher or ↓lower). The abbreviations list and explanations of the cardiovascular parameters are included in Supplemental Table 1.

**Supplemental Table 8.** Transcriptomic analysis of *fbn2b^-/-^* mutants.

(Tab 1 and 2) List of all differentially expressed genes (DEG) in *fbn2b^-/-^* mutants to WT control siblings at 1 and 2 dpf, based on logFC and adjusted p-value (FDR). (Tab 3 and 4) Overview of all gene ontology-enriched biological pathways in *fbn2b^-/-^* mutants to WT control siblings at 1 and 2 dpf. Supplemental Table 8 is attached as a separate file.

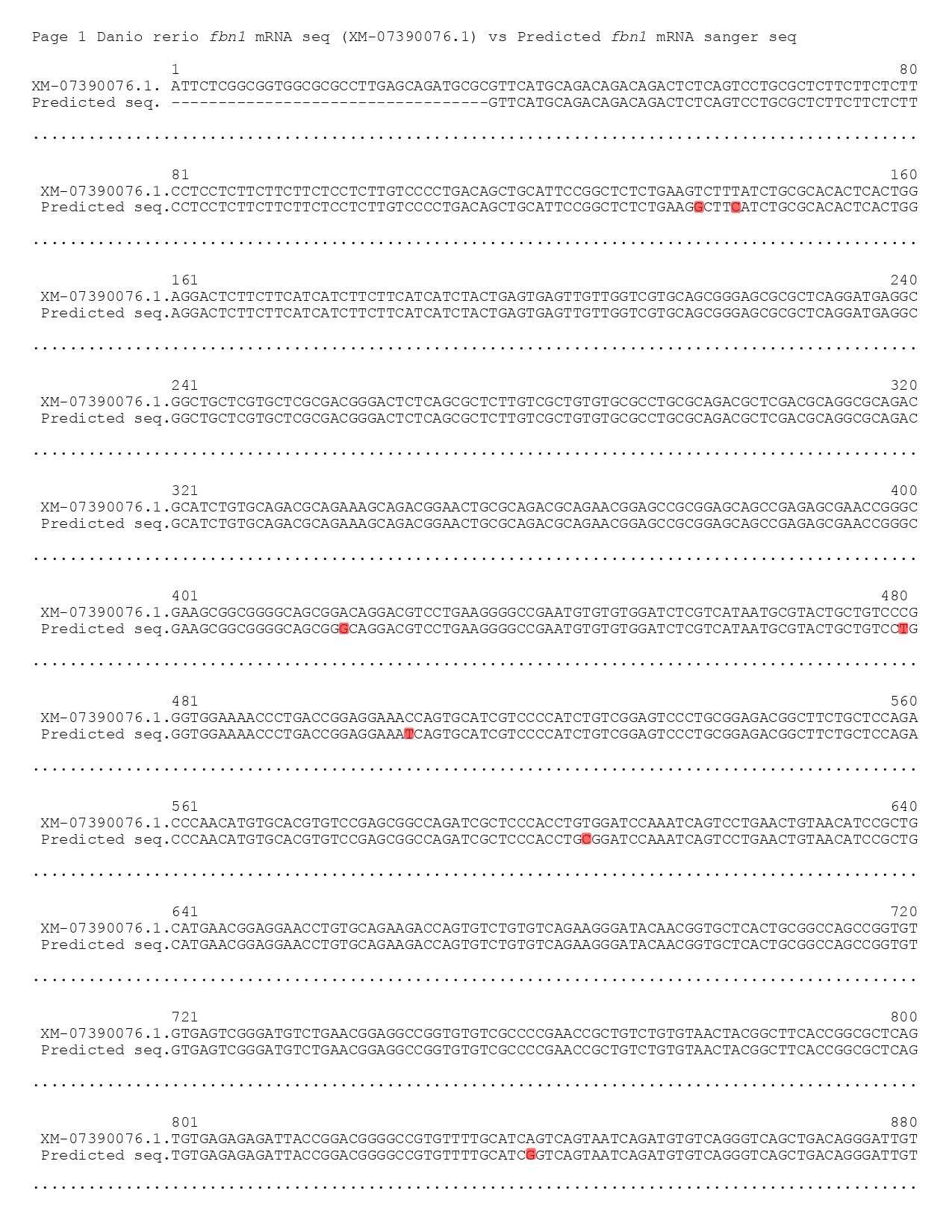

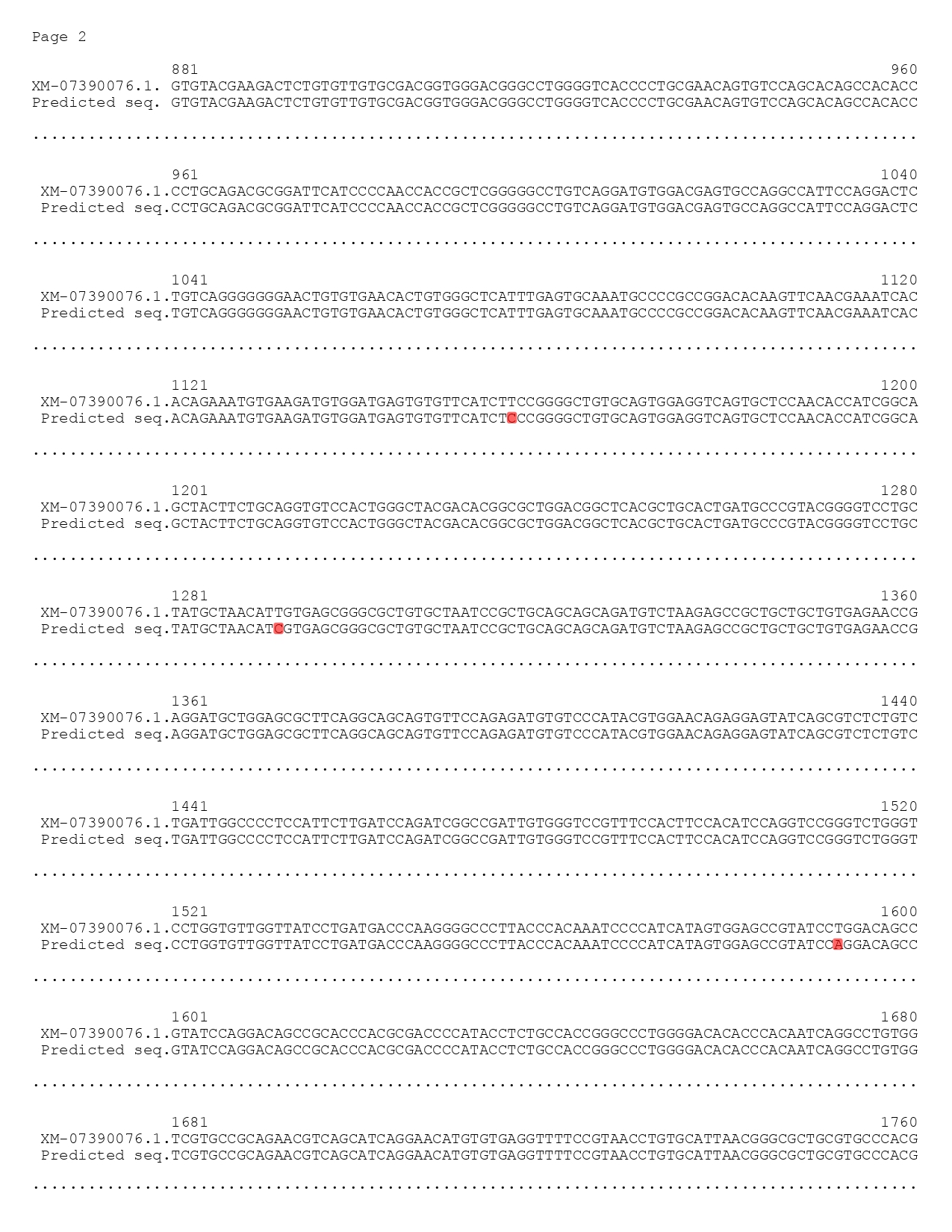

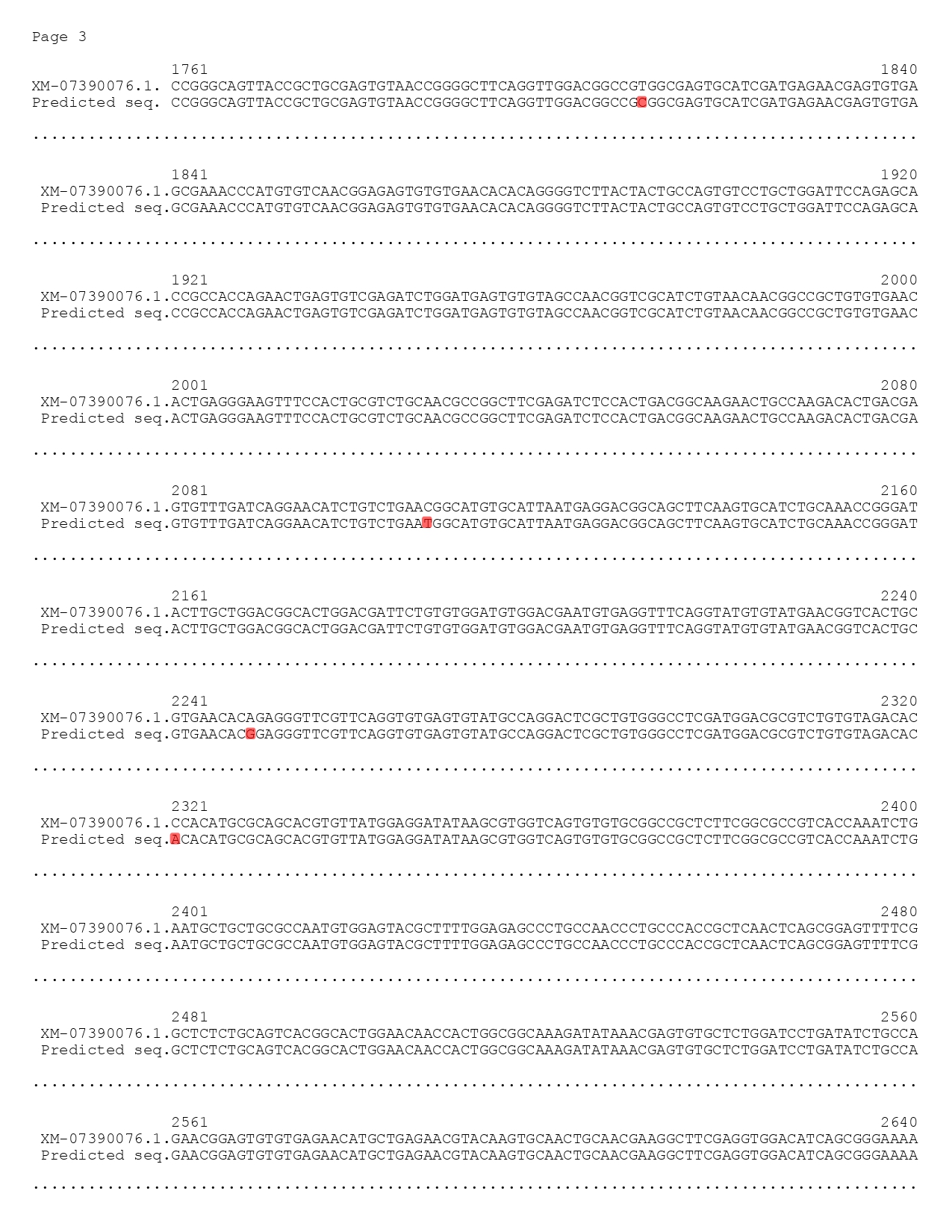

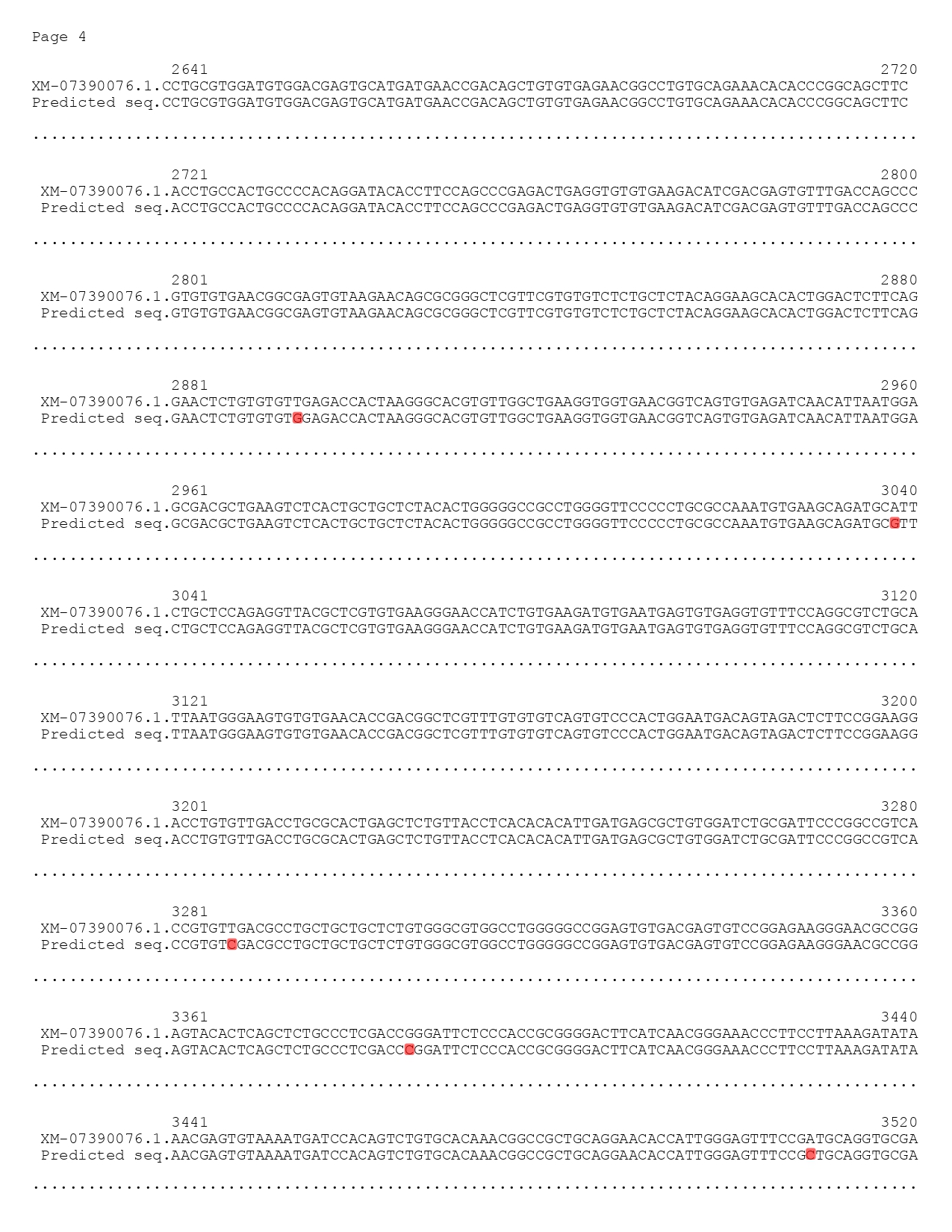

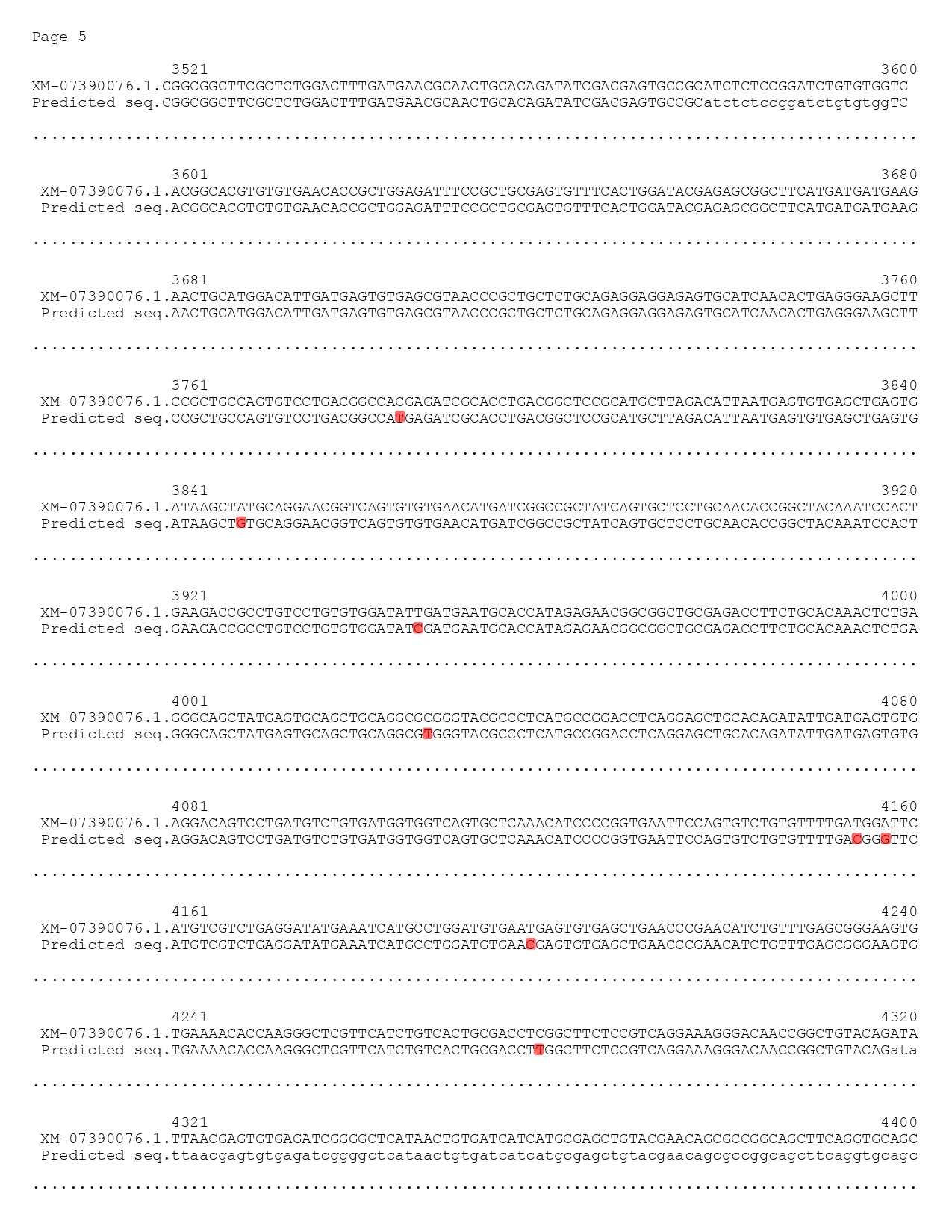

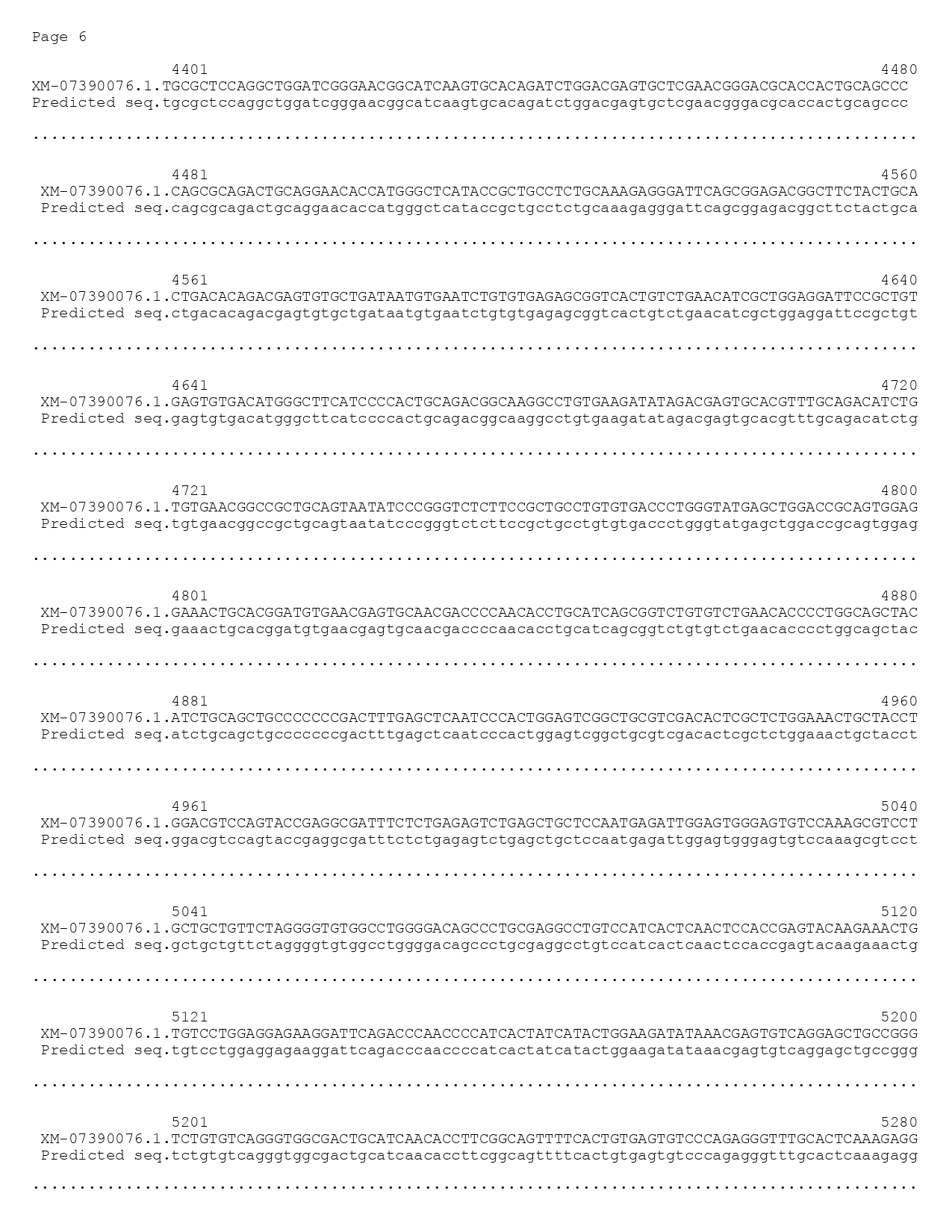

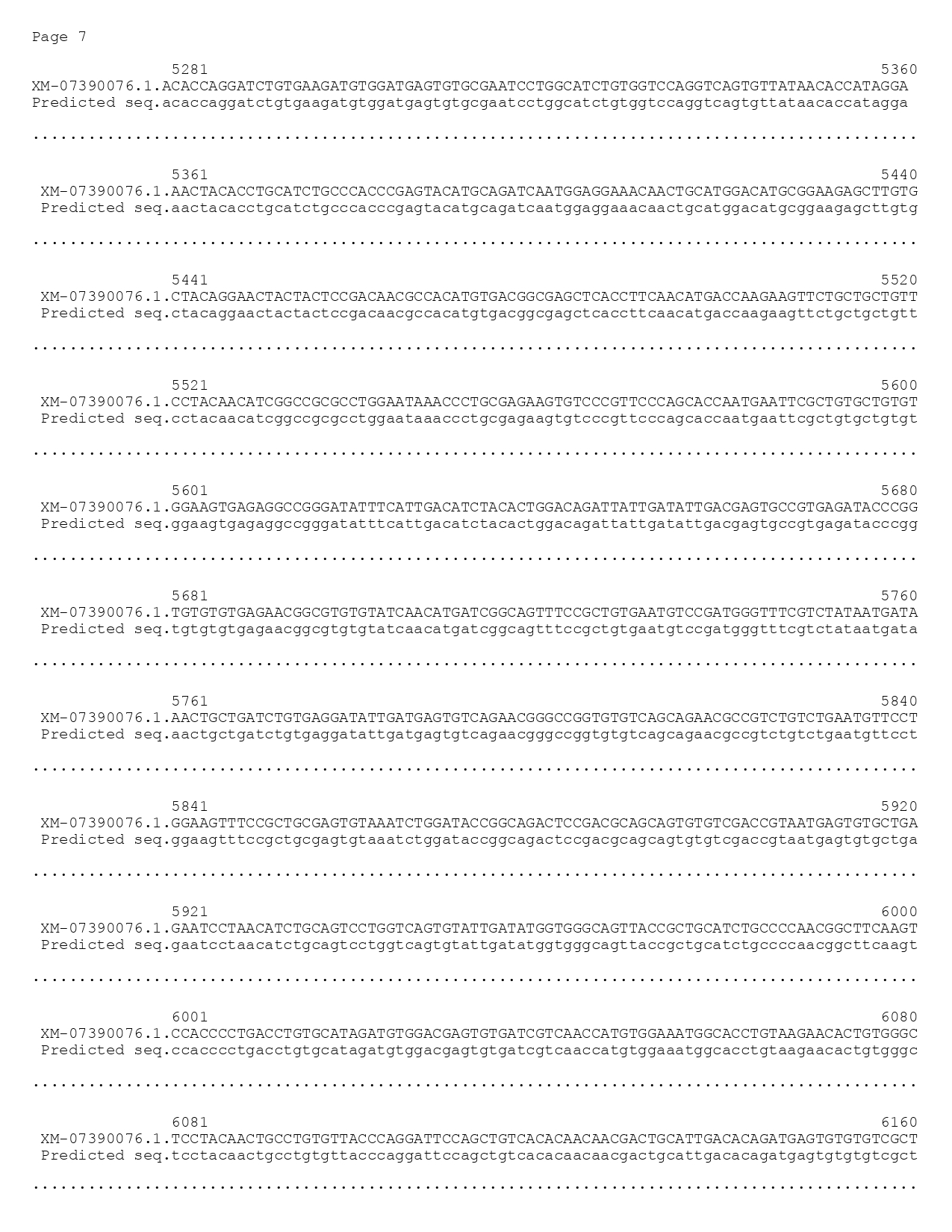

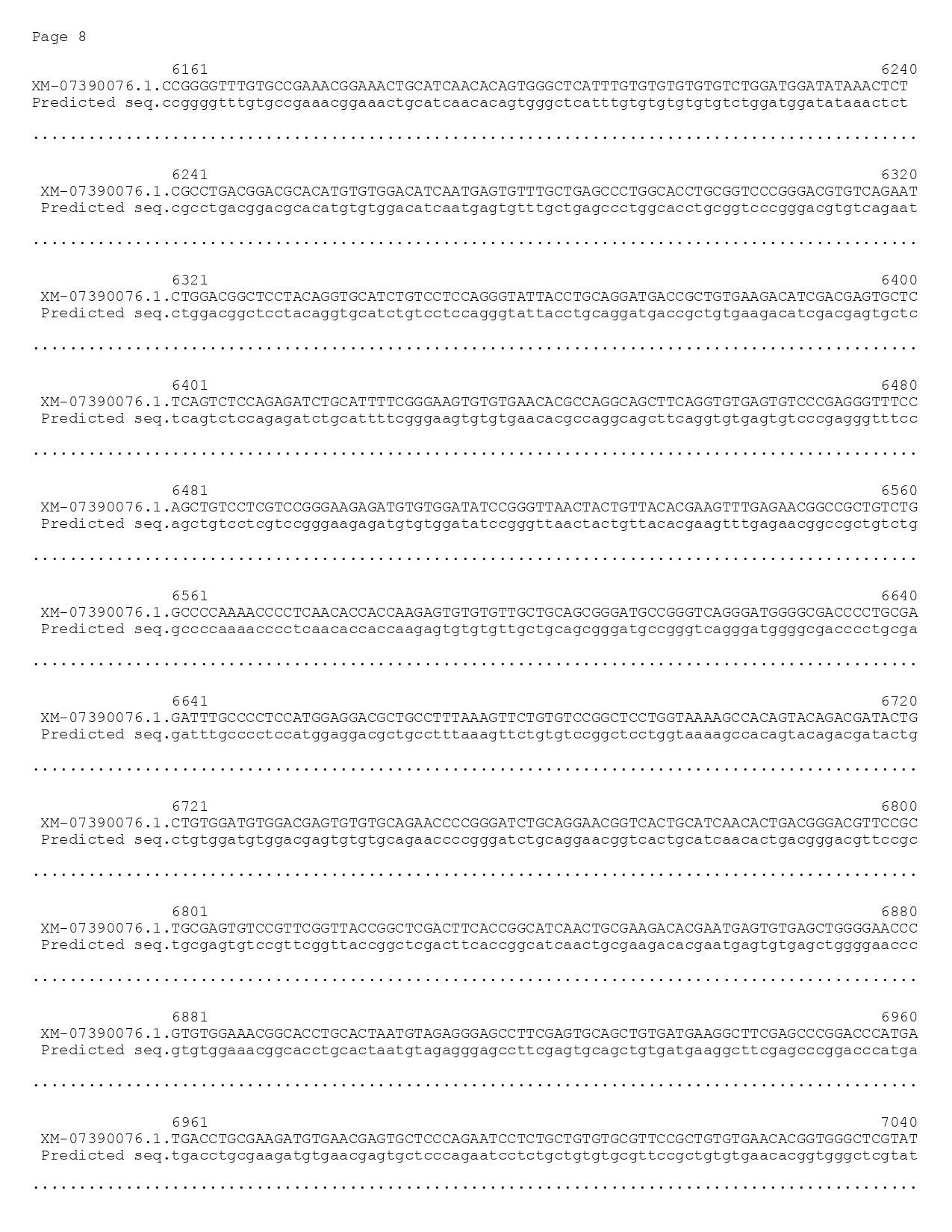

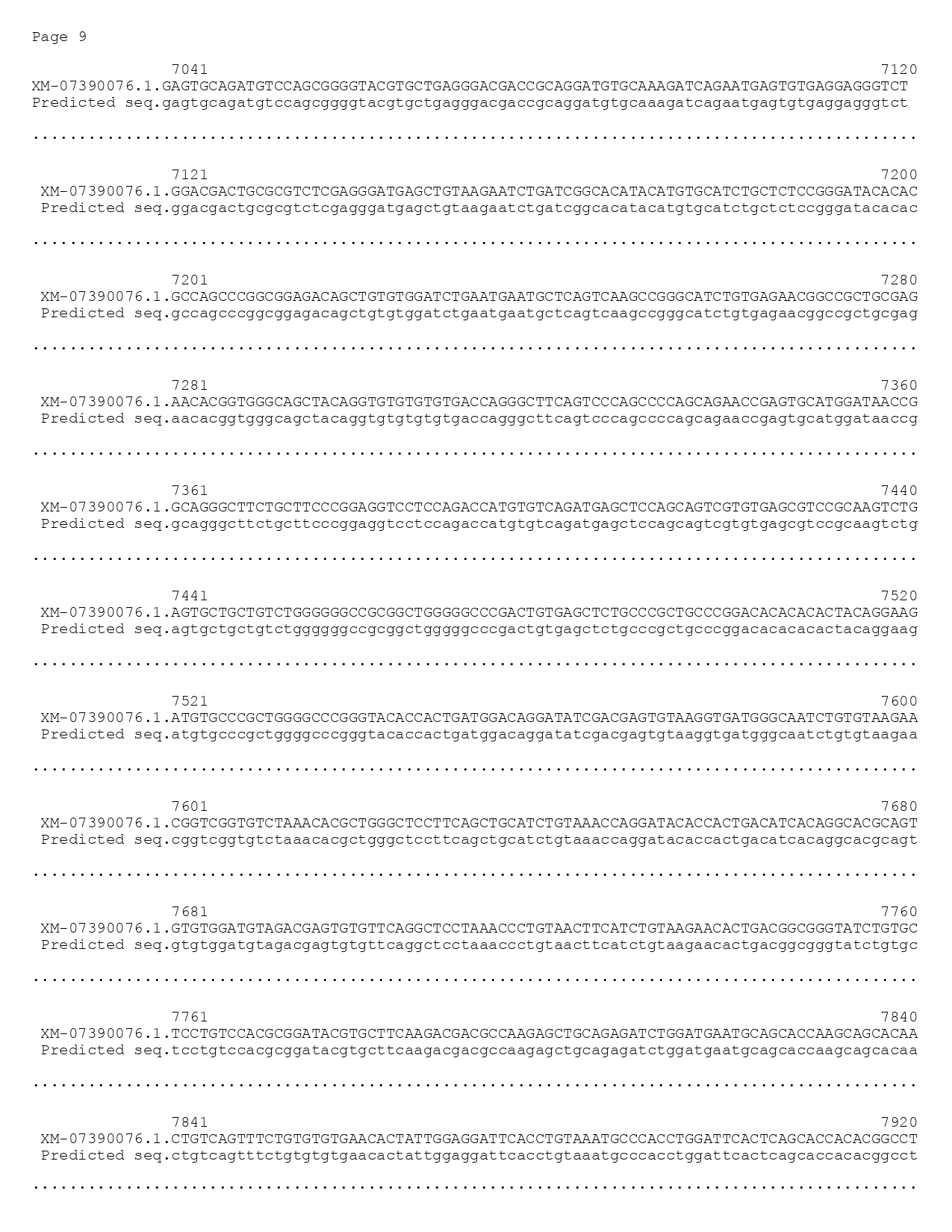

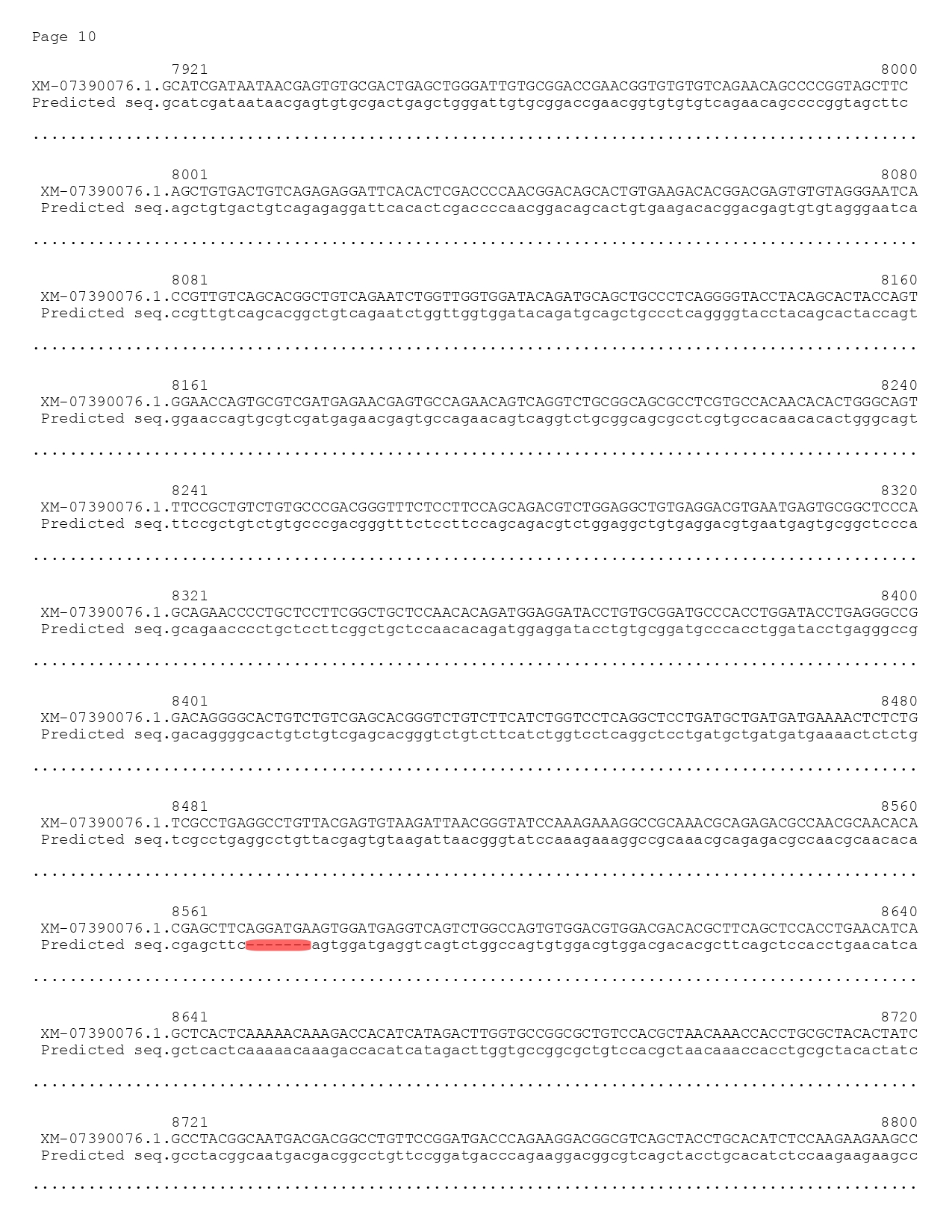

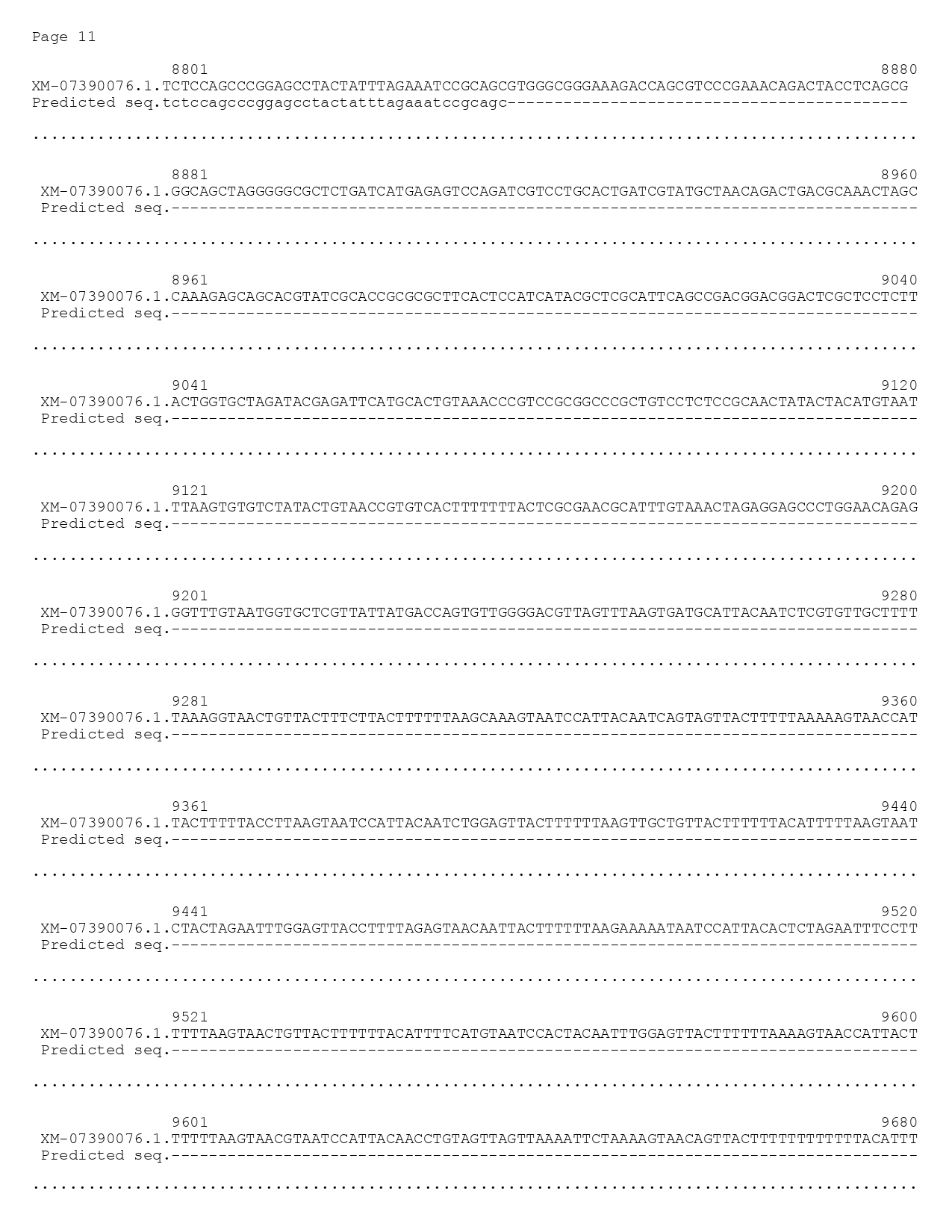

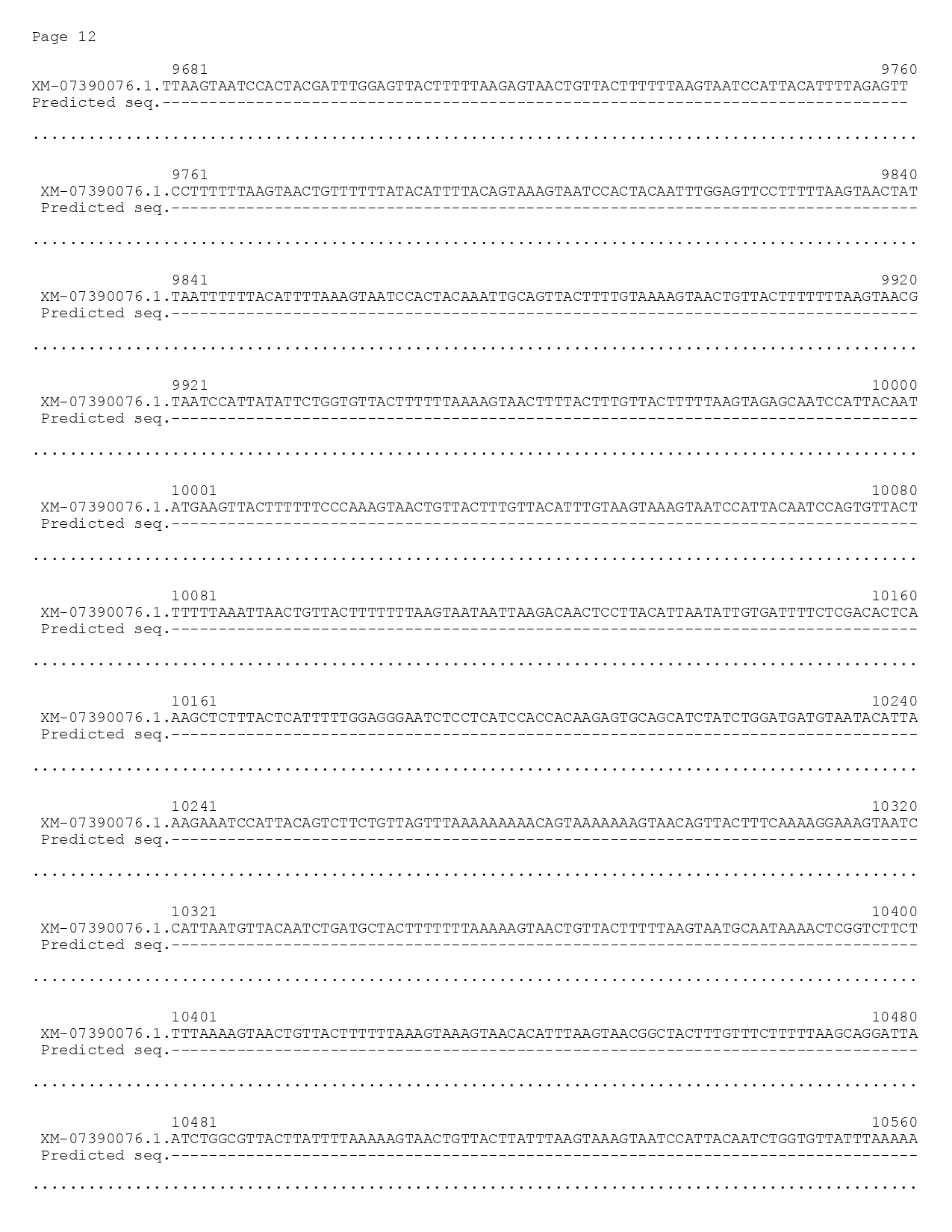

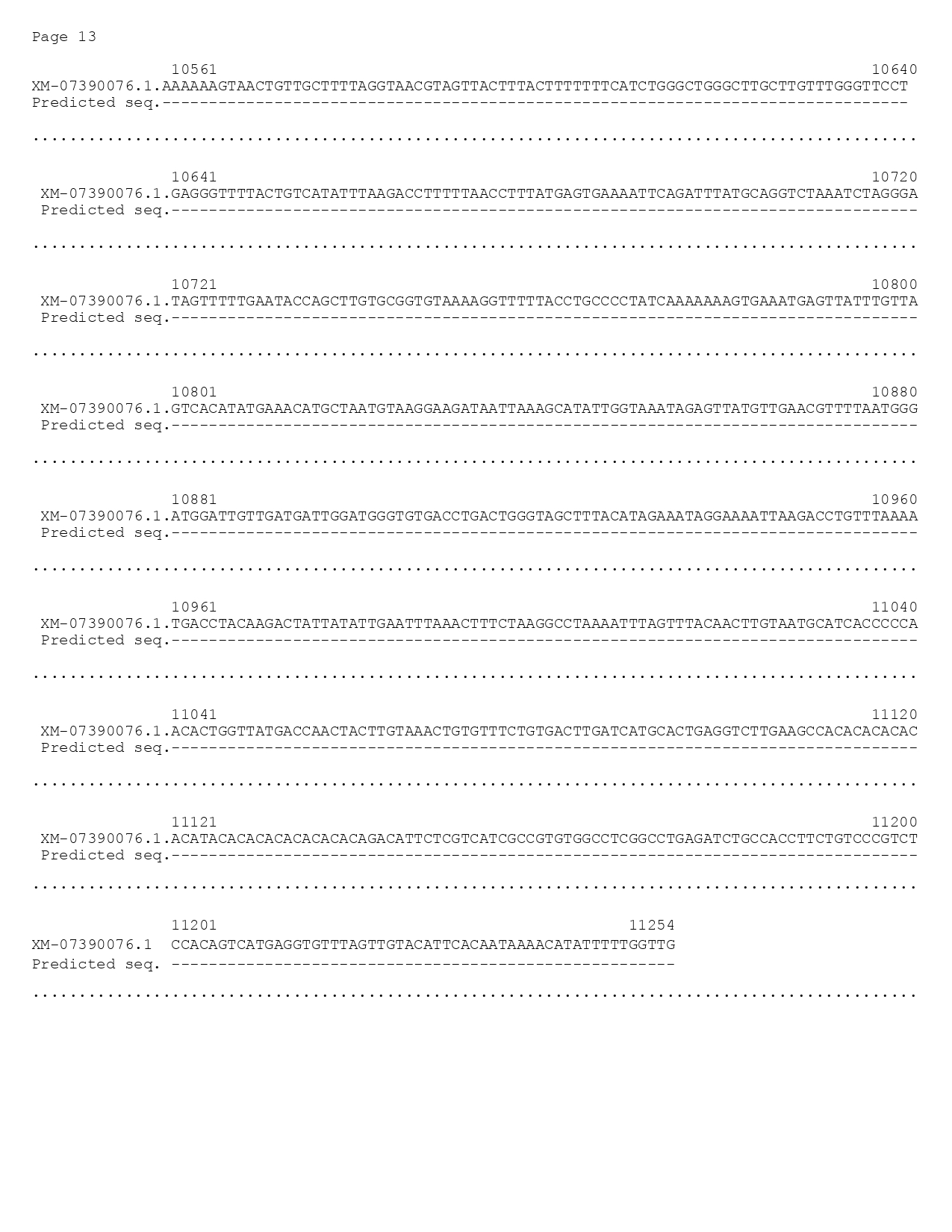

**Supplemental Figure 1.** *Prediction of full lenghth fbn1 mRNA sequence (XM_073930076.1).*

Alignment of the *Danio rerio* *fbn1* mRNA sequence (XM_073930076.1) to our in-house predicted mRNA fbn1 sequence through sanger sequencing. Red indicates single nucleotide polymorphisms (SNP’s).

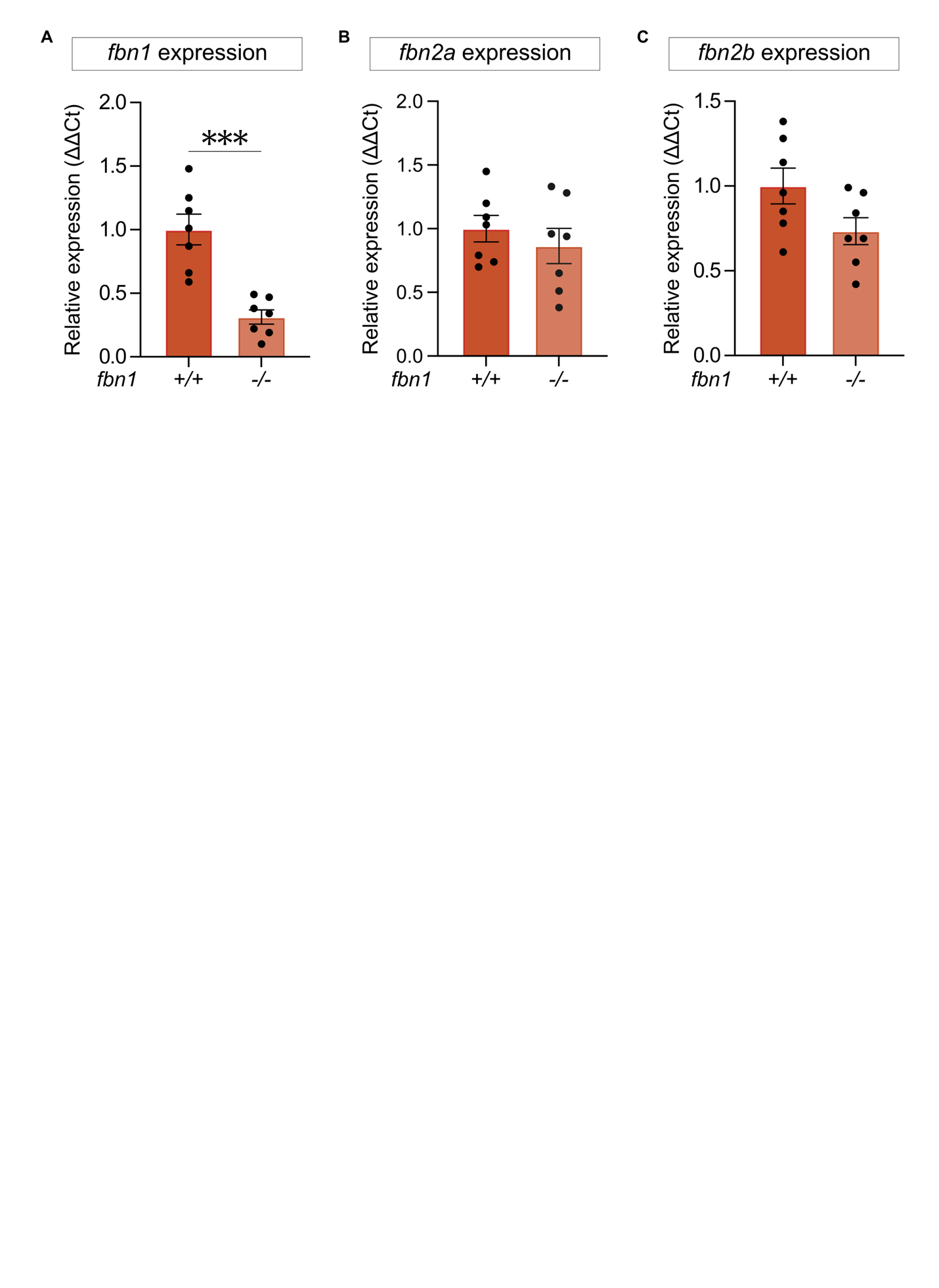

**Supplemental Figure 2.** *fbn1*, *fbn2a* and *fbn2b* expression in fbn1^-/-^ zebrafish.

RT-qPCR analysis of *fbn1* **(A)**, *fbn2a* **(B)** and *fbn2b* **(C)** mRNA expression in 5 dpf WT and *fbn1^-/-^* (Cmg80) larvae (n = 7). Each data points represents the mean of 2 technical repeats. Data are expressed as mean ± SEM. Statistical analysis: unpaired t-test. ***p<0.001.

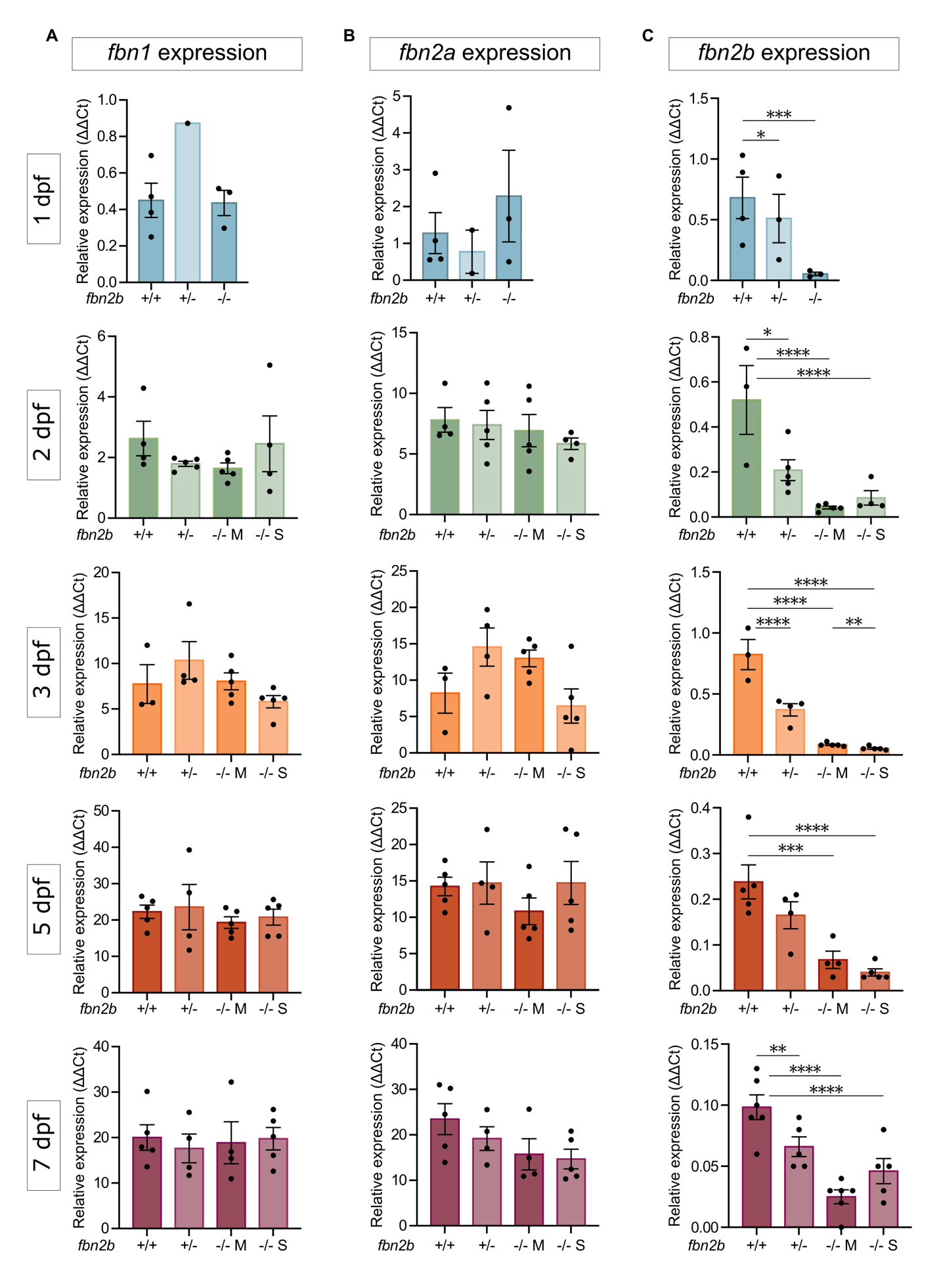

**Supplemental Figure 3.** *fbn1*, *fbn2a* and *fbn2b* expression patterns in *fbn2b^-/-^* zebrafish.

RT-qPCR analysis of *fbn1* **(A)**, *fbn2a* **(B)** and *fbn2b* **(C)** mRNA expression in WT and *fbn2b^-/-^* (mild and severe) on RNA extracted from whole embryos at 1, 2, 3, 5 and 7 dpf (n = 2-5 for each developmental stage). Each datapoint represents the mean of two technical replicates. Statistical analysis: One-way ANOVA followed by Tukey multiple comparison’s test on log-transformed data and tested for trend. Data are expressed as mean ± SEM. ****p<0.0001, ***p<0.001, **p<0.01, *p<0.05., M = mild pericardial phenotype, S = severe pericardial phenotype.

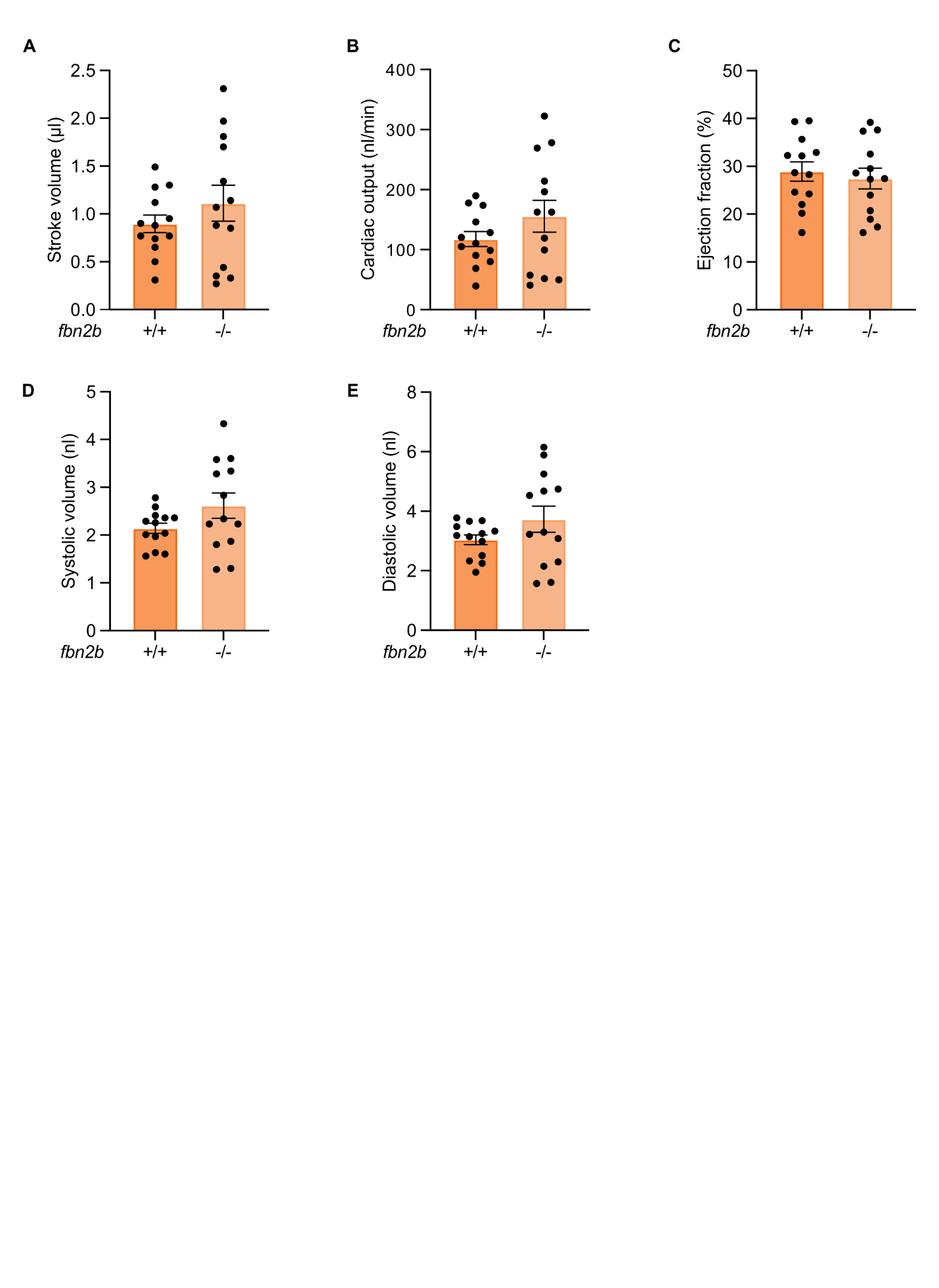

**Supplemental Figure 4.** Heart function analysis of 3 dpf *fbn2b^-/-^* zebrafish.

Brightfield microscopy was used to quantify cardiac parameters in 3dpf *fbn2b^-/-^* larvae and WT siblings: **(A)** stroke volume (mL), **(B)** cardiac output (nl/min), **(C)** ejection fraction (%), **(D)** systolic volume (Vsys, nl) and **(D)** diastolic volume (Vdias, nl) ( n = 13). Data are expressed as a mean ± SEM. Statistical test analysis: unpaired t-test. No statistical difference was reached.

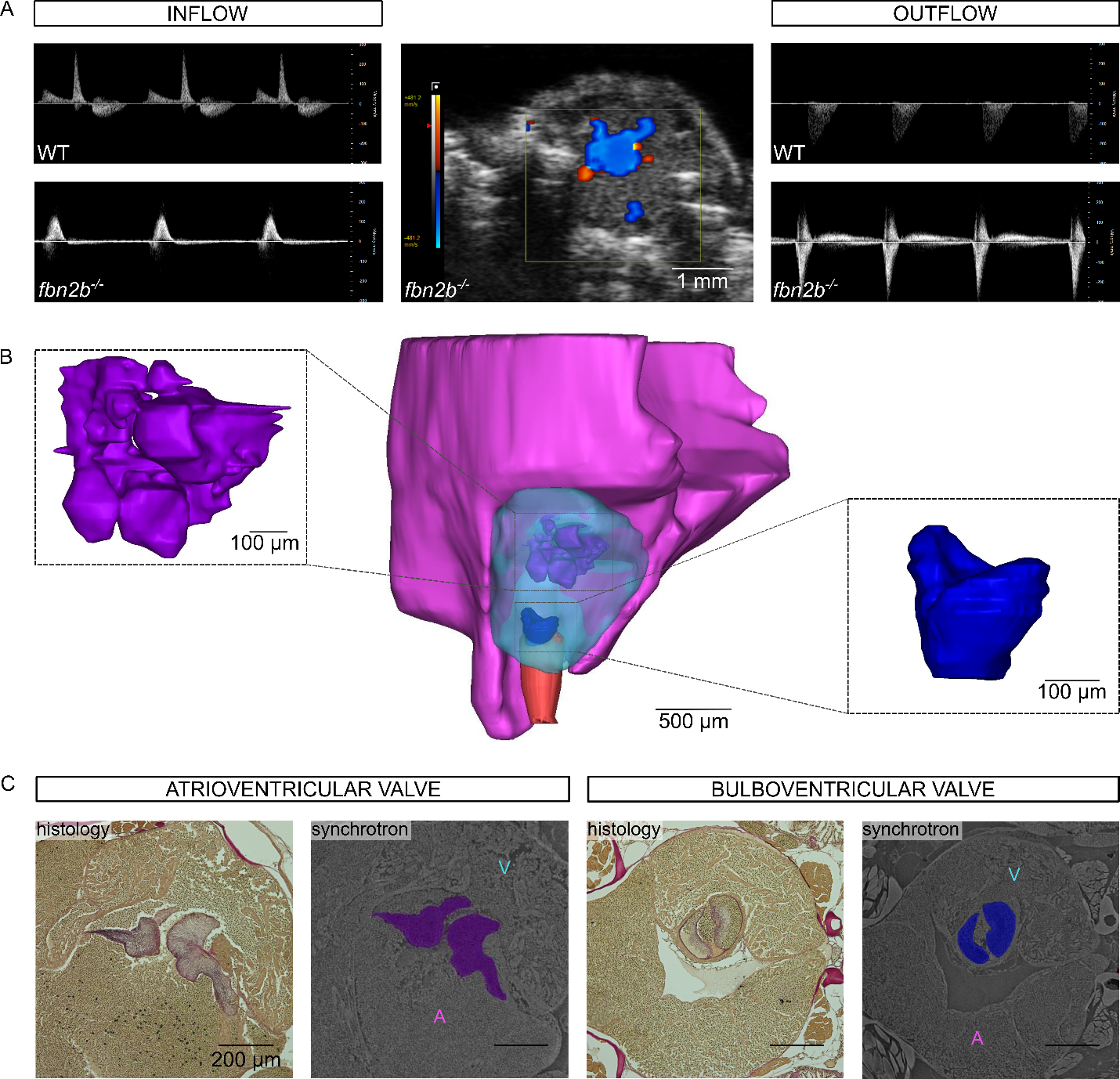

**Supplemental Figure 5.** Multimodal assessment of a *fbn2b^-/-^* zebrafish with an exceptionally severe cardiac phenotype.

(A) Pulse Wave Doppler images showing blood inflow into the ventricle (left) and outflow from the ventricle into BA (right), of WT zebrafish (top) in comparison to a highly abnormal *fbn2b^-/-^* zebrafish (bottom). This mutant zebrafish exhibited pronounced regurgitation of the BV valve, which is evident in CFD imaging, showing simultaneous inflow (orange) and outflow (blue) at the same location (middle image). (B) 3D heart model of the same mutant zebrafish, created by synchrotron X-ray scanning. The atrium is displayed in pink, the ventricle in slightly translucent blue, BA in red, the AV valve in purple, and the BV valve in dark blue. The AV valve appears amorphic with various folded and fused segments, as well as being highly hypermorphic. The atrium is notably large, so much so that its uppermost segment was not captured in the synchrotron field of view. In contrast, the ventricle and BA are relatively small. (C) The same AV and BV valve sections were visualised using histological elastin staining (purple) and synchrotron imaging (purple and blue), demonstrating the similarities between the two techniques. A = atrium, V = ventricle, WT = wild type.

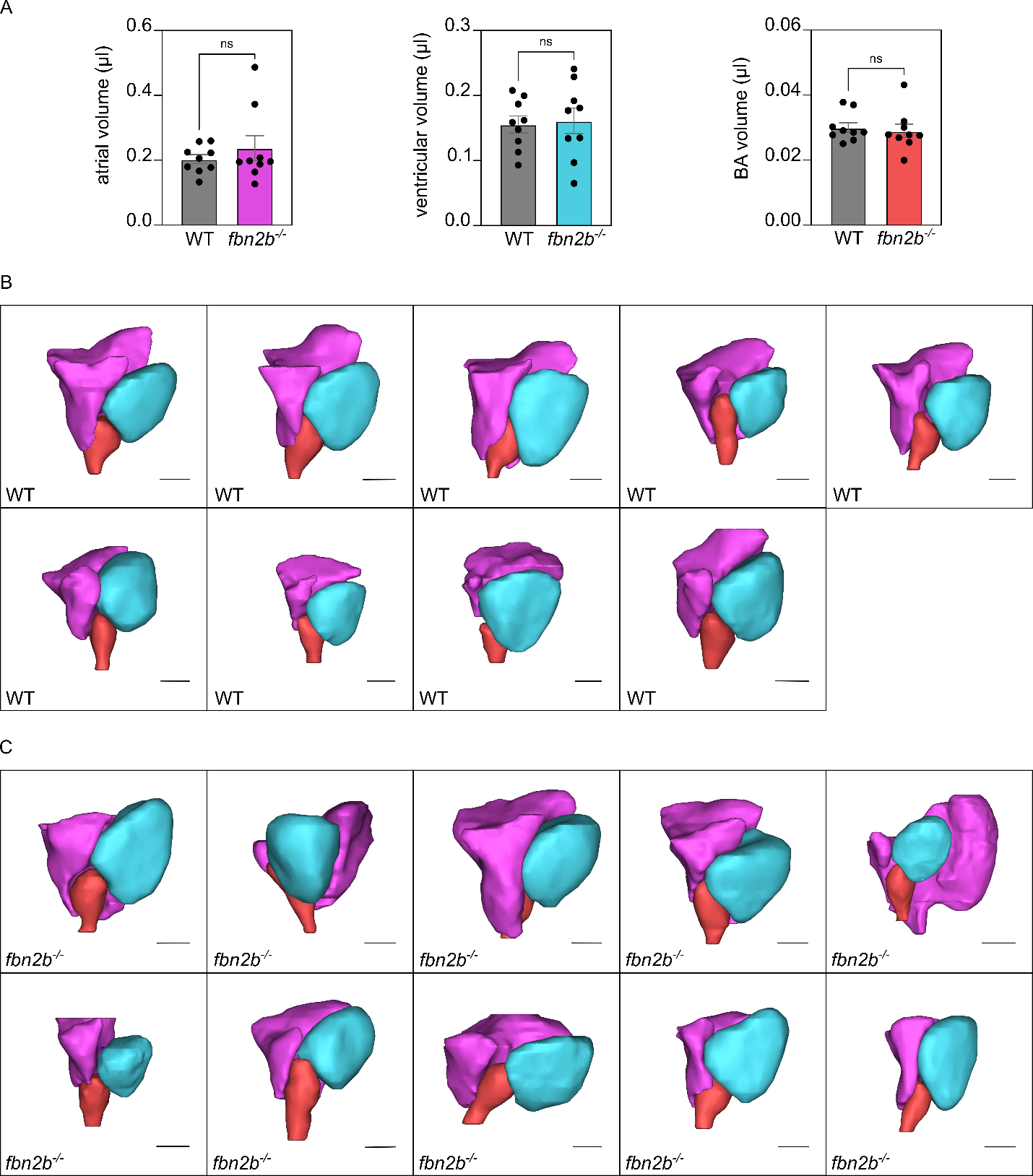

**Supplemental Figure 6.** 3D reconstructions of whole zebrafish hearts.

(A) Volumes of atrium, ventricle, and BA were obtained from 3D models based on synchrotron X-ray imaging. Reported volumes include combined lumen and wall. While no statistically significant differences were observed between WT and *fbn2b^-/-^* zebrafish, a greater variability was evident in the mutant group. These differences are visually apparent in the 3D zebrafish heart models of WT (B) or *fbn2b^-/-^* mutants (C). The atrium is displayed in pink, the ventricle in slightly translucent blue, and BA in red. Data are expressed as a mean ± SEM. ns = non-significant. Statistical test analysis: unpaired t-test. Scale bar: 500 µm. WT = wild type.

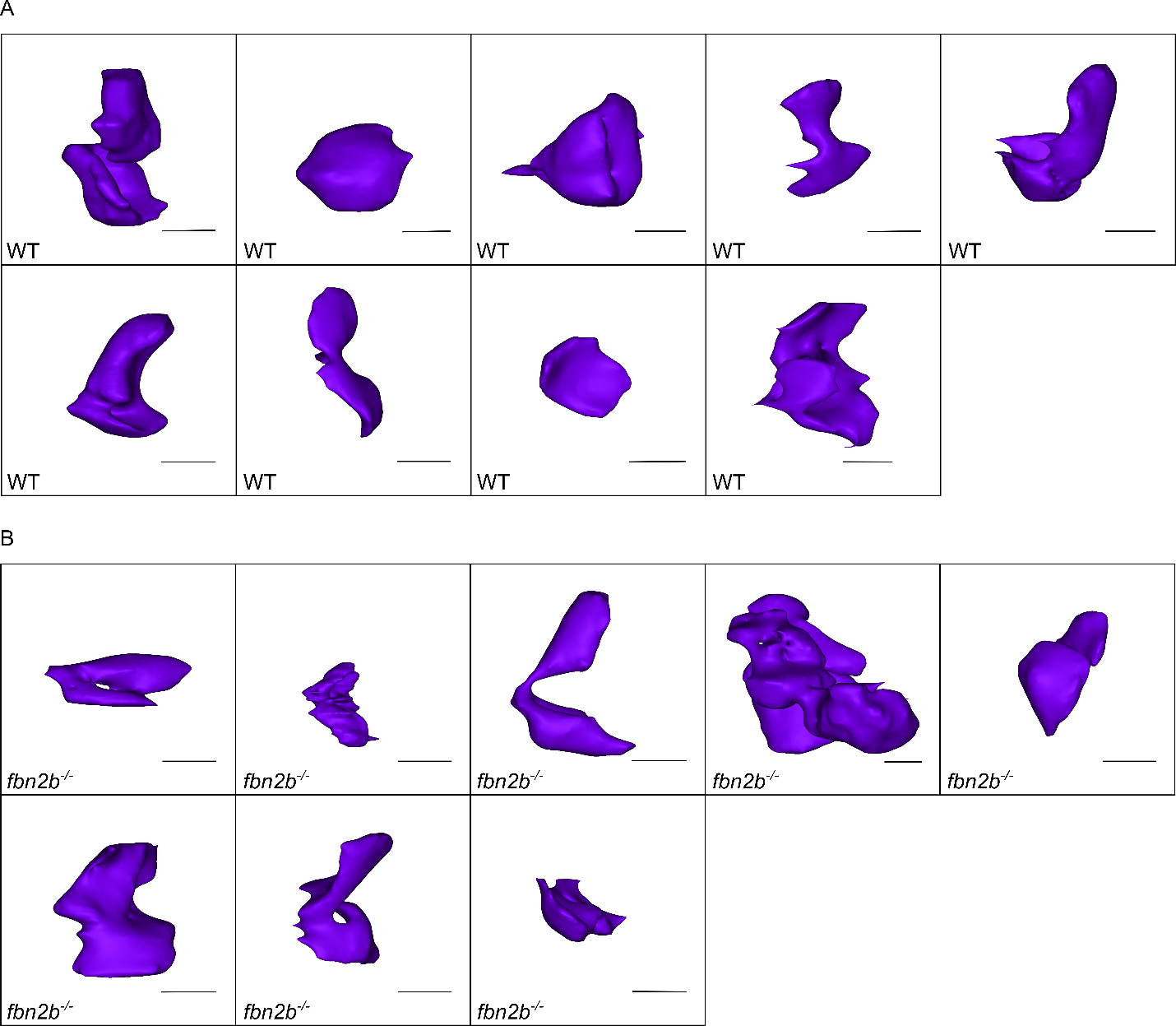

**Supplemental Figure 7.** 3D reconstructions of zebrafish AV valves.

Reconstructions based on synchrotron scans of AV valves from WT (A) and *fbn2b^-/-^* mutants (B). The shape of the valve leaflets varied depending on their position during sample fixation, leading to inconsistencies. As a result, precise 2D dimensions (e.g. length and thickness) could not be accurately determined. All images from the 3D models were captured from the same perspective. Scale bar: 100 µm. WT = wild type.

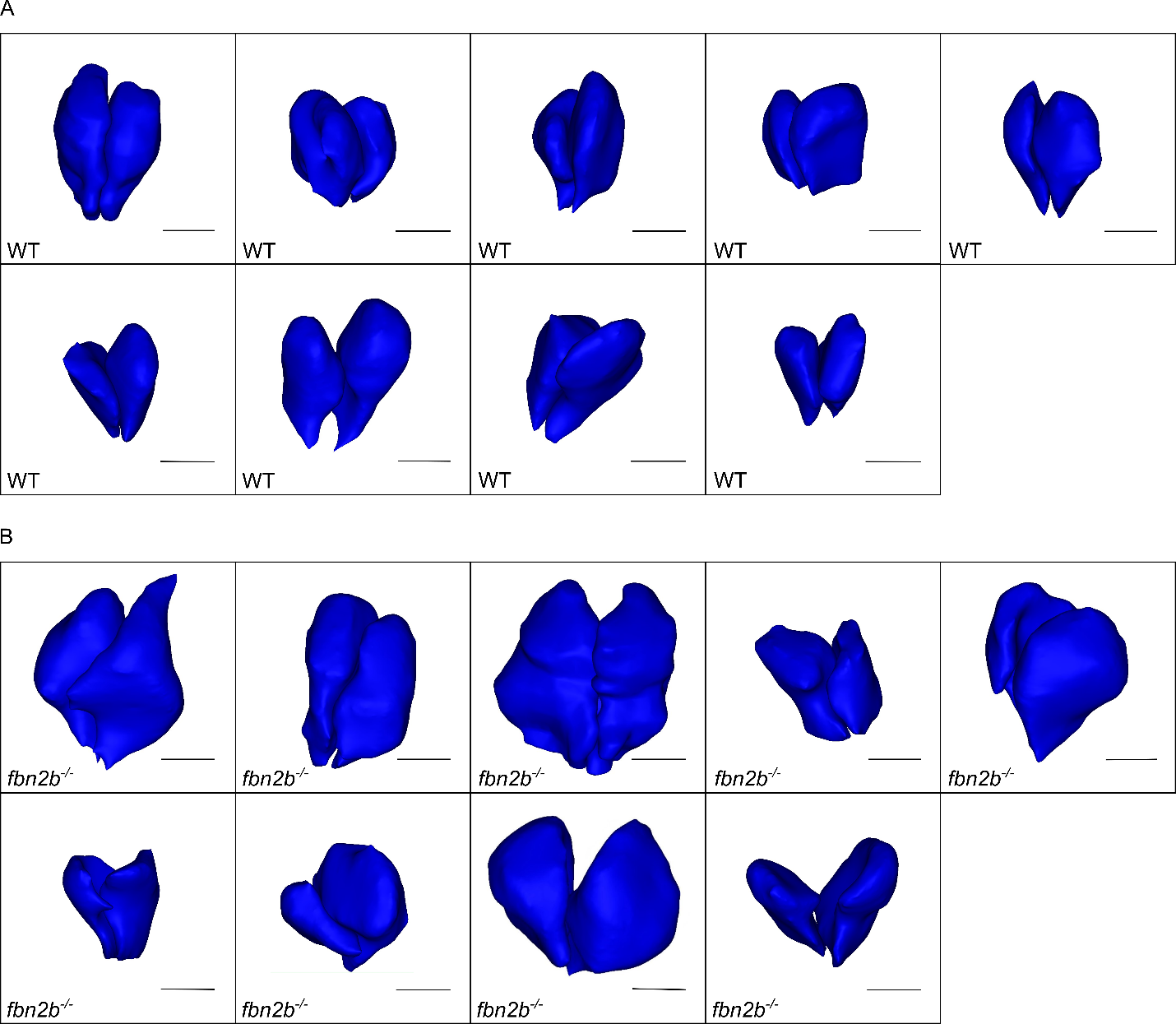

**Supplemental Figure 8.** 3D reconstructions of zebrafish BV valves.

Reconstructions based on synchrotron scans of BV valves from WT (A) and *fbn2b^-/-^* mutants (B). The difference in leaflet sizes between WT and mutants can be visually observed. All images from the 3D models were captured from the same perspective. Scale bar: 100 µm. WT = wild type.

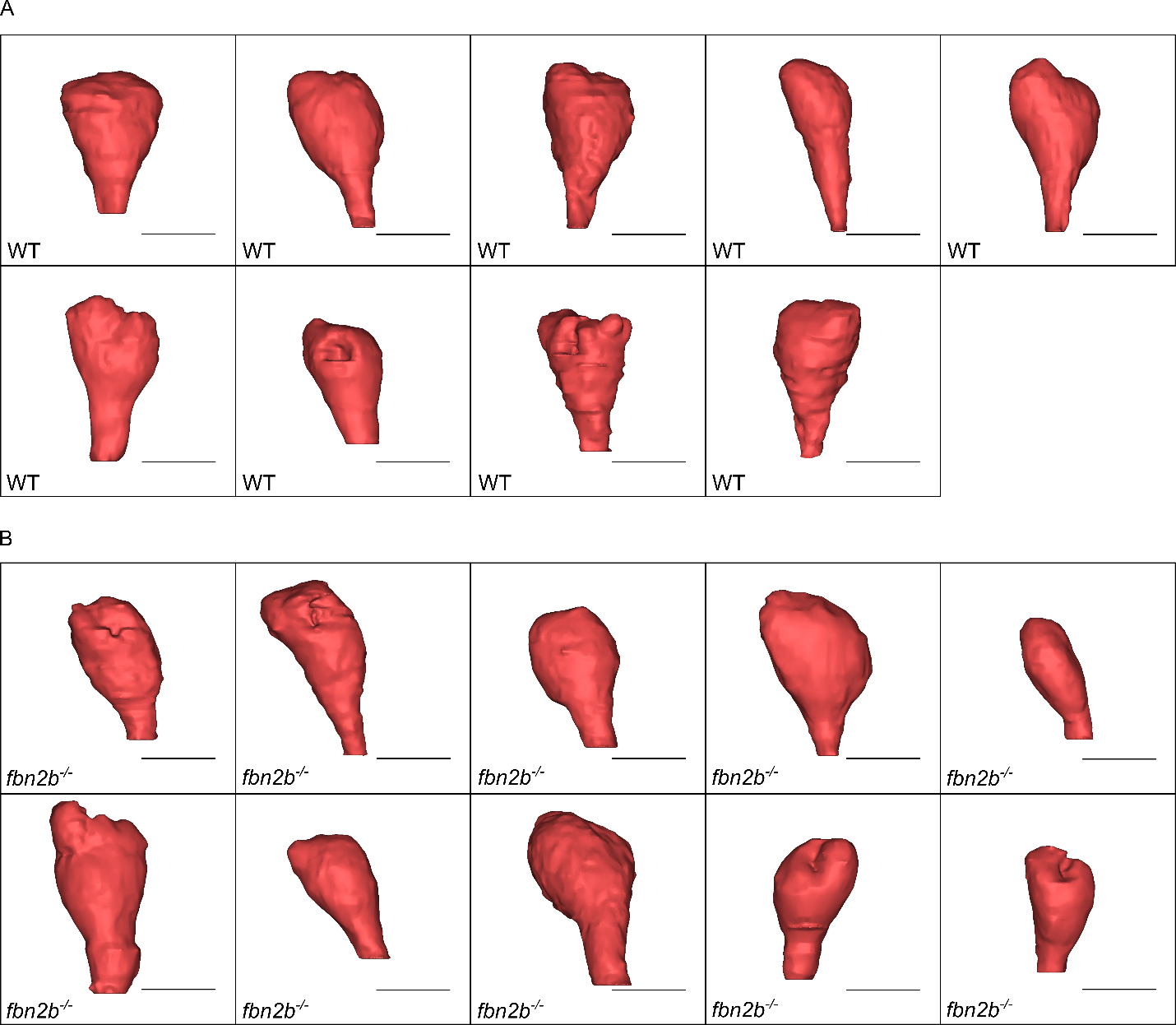

**Supplemental Figure 9.** 3D reconstructions of zebrafish BA.

Reconstructions based on synchrotron scans of BA from WT **(A)** and *fbn2b^-/-^* mutants **(B)**. All images from the 3D models were captured from the same perspective. Scale bar: 100 µm. WT = wild type.

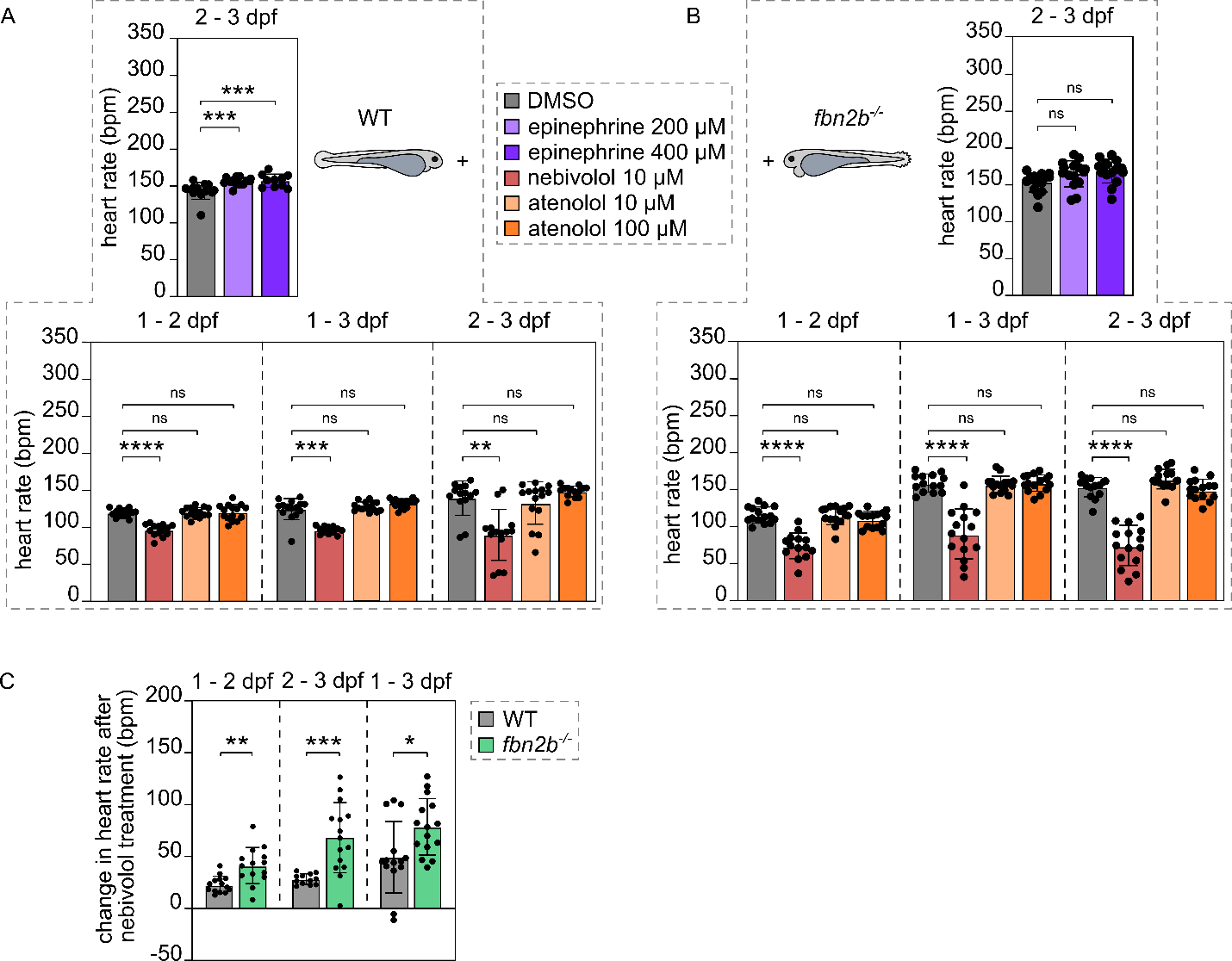

**Supplemental Figure 10.** Heart rate modulation by β-adrenergic receptor agonists and antagonists in zebrafish embryos.

**(A)** WT and **(B)** *fbn2b^-/-^* zebrafish embryos were treated with β-adrenergic receptor antagonists (nebivolol 10 µM – red, atenolol 10 µM and 100 µM – orange) and β-adrenergic receptor agonist (epinephrine 200 µM and 400 µM – purple), with solvent as a control (DMSO – grey) (n = 12 – 15). Heart rate was measured in beats per minute (bpm) and compared with the controls. (C) The change in heart rate after nebivolol (1 µM) treatment in WT (grey) and *fbn2b^-/-^* zebrafish (green), measured as the difference between the mean heart rate in untreated controls and the heart rate in the treated zebrafish, for each sample individually. Data are expressed as a mean ± SEM. ****p<0.0001, ***p<0.001, **p<0.01, ns = non-significant. Statistical test analysis: one-way ANOVA with Dunnett’s multiple comparisons test (A top, bottom 1 – 2 dpf; B bottom), Kruskal-Wallis test with Dunn’s multiple comparisons test (A bottom 1 – 3 dpf, 2 – 3 dpf; B top), unpaired t-test (C). WT = wild typ
